# Sensory neuron fate is developmentally perturbed by *Gars* mutations causing human neuropathy

**DOI:** 10.1101/071159

**Authors:** James N. Sleigh, John M. Dawes, Steven J. West, Emily L. Spaulding, Adriana Gómez-Martín, Robert W. Burgess, M. Zameel Cader, Kevin Talbot, David L. Bennett, Giampietro Schiavo

**Author notes:** ^#^ These authors contributed equally (John M. Dawes, Steven J. West). Correspondence to: James N. Sleigh, Giampietro Schiavo.

## Abstract

Charcot-Marie-Tooth disease type 2D (CMT2D) is a peripheral nerve disorder caused by dominant, toxic, gain-of-function mutations in the widely expressed, housekeeping gene, *GARS*. The mechanisms underlying selective nerve pathology in CMT2D remain unresolved, as does the cause of the mild-to-moderate sensory involvement that distinguishes CMT2D from the allelic disorder distal spinal muscular atrophy type V. To elucidate the mechanism responsible for the underlying afferent nerve pathology, we examined the sensory nervous system in CMT2D mice. We show that the equilibrium between functional subtypes of sensory neuron in dorsal root ganglia is distorted by *Gars* mutations, leading to sensory defects in peripheral tissues and correlating with overall disease severity. CMT2D mice display changes in sensory behaviour concordant with the afferent imbalance, which is present at birth and non-progressive, indicating that sensory neuron identity is prenatally perturbed and that a critical developmental insult is key to the afferent pathology. This suggests that both neurodevelopmental and neurodegenerative mechanisms contribute to CMT2D pathogenesis, and thus has profound implications for the timing of future therapeutic treatments.

**Significance Statement:** Charcot-Marie-Tooth disease (CMT) is a collection of genetically diverse inherited nerve disorders with the unifying feature of peripheral neuron degeneration. The mechanisms triggering this motor and sensory nerve dysfunction remain unresolved, as does the reason for the lack of sensory pathology observed in distal hereditary motor neuropathies, which can be associated with CMT genes. To unravel the mechanisms leading to afferent deterioration, we have studied the sensory nervous system of CMT Type 2D mice. Our work indicates that the specific cellular identity of sensory nerves is perturbed in mutant mice pre-natally. CMT therefore manifests through the complex interplay between malfunctioning developmental, maturation, and survival programs, which has important ramifications for therapeutic timing.

## Introduction

Charcot-Marie-Tooth disease (CMT) is a group of genetically diverse peripheral neuropathies that share the main pathological feature of progressive motor and sensory degeneration (1). Although lifespan is usually unaffected, patients display characteristic muscle weakness and wasting predominantly in the extremities, leading to difficulty walking, foot deformities, and reduced dexterity (2). CMT is traditionally divided into type 1 or demyelinating CMTs that display loss of peripheral nerve myelin causing reduced nerve conduction velocity (NCV), type 2 or axonal CMTs typified by axon loss with relatively normal NCVs, and intermediate CMTs that share clinical features of CMT1 and CMT2 (1). Over 80 different genetic loci have been linked to CMT, which is known to affect ≈1/2,500 people, making it the most common group of hereditary neuromuscular disorders (3).

Dominant mutations in the glycyl-tRNA synthetase (GlyRS) gene, *GARS*, are causative of CMT type 2D (CMT2D, OMIM 601472), which normally manifests during adolescence and presents with muscle weakness in the upper extremities, followed by the feet (4). The 2D subtype is one of a number of CMTs associated with mutation of an aminoacyl-tRNA synthetase (ARS) gene (5-8). Humans possess 37 ARS proteins, which covalently link amino acids to their partner transfer RNAs (tRNAs), thereby charging and priming the tRNAs for protein synthesis. This housekeeping function of glycine aminoacylation explains the widespread and constitutive nature of *GARS* expression (4), but highlights the phenomenon of neuronal specificity in the disease: why do mutations that affect a ubiquitous protein selectively trigger peripheral nerve degeneration? Several hypotheses have been suggested (9, 10), although the exact disease mechanisms remain unknown. Nevertheless, cell-based experiments and studies using two CMT2D mouse models (the mild *Gars*^*C201R*/+^ allele and the more severe *Gars*^*Nmf249*/+^ model) indicate that CMT2D is likely caused by a toxic gain-of-function in mutant GlyRS rather than haploinsufficiency or a loss of aminoacylation activity or a non-canonical function (11-15). A possible mediator of this toxicity was identified when five CMT2D-associated mutations spread along the length of *GARS* were all shown to induce a similar conformational change in GlyRS, leading to the exposure of surfaces buried in the wild-type protein (16). These neomorphic regions likely facilitate the aberrant accumulation of mutant GlyRS at the neuromuscular junction (NMJ) of a CMT2D *Drosophila melanogaster* model (17), and the erroneous interaction of mutant GlyRS with NRP1, antagonising VEGF signaling (18).

A second major conundrum in GlyRS-associated neuropathy is why some patients with dominant *GARS* mutations, diagnosed with the allelic neuropathy distal spinal muscular atrophy type V (dSMA-V, OMIM 600794) (4), lack the distinguishing mild-to-moderate sensory involvement typical of CMT2D (19-22). The ability of CMT2D patients to sense vibration is most impaired, followed by light touch, temperature, and pain (19). Furthermore, CMT2D patients display deficits in deep tendon reflexes of the extremities (21, 22), while reflexes of dSMA-V patients remain relatively unperturbed (4, 23), implicating defective relay arc afferents rather than efferents. CMT2D sensory defects are dependent on disease severity not duration, while dSMA-V patients are refractory to sensory pathogenesis, suggesting that the two disorders lie along a spectrum and that disease-modifying loci may dictate these differences (19). Accordingly, CMT2D and dSMA-V can be caused by the same *GARS* mutation and manifest at different ages within a family (20).

CMT2D sensory pathology, both in patients and animal models, has not been studied in detail, although the limited sensory analysis has identified possible contradictions that require clarification. The greatest sensory deficiency in CMT2D patients is in the perception of vibration, which is sensed by neurons with large cell bodies and axons (24, 25); however, patient sural nerve biopsies show a selective loss of small sensory axons (19, 20). This histological finding is also counter to what is observed in CMT2D mice; the milder *Gars*^*C201R*/+^ mice display a general reduction in axon diameter in both the saphenous and sensory femoral nerves (12), while the more severe *Gars*^*Nmf249*/+^ allele displays both a reduction in axon diameter and axon number (11); nevertheless, whether specific sensory neuron populations are preferentially atrophied or lost is unknown. We thus set out to interrogate the sensory nervous system of CMT2D mice to better understand how and when *Gars* mutations cause sensory pathology, and to determine the effect that this has on sensation of the external environment.

## Results

### Gars^C201R/+^ dorsal root ganglion (DRG) cultures have a smaller percentage of large area sensory neurons

We began our CMT2D sensory nervous system analysis by culturing primary DRG neurons from wild-type and *Gars*^*C201R*/+^ mice. This model of CMT2D has a mutagen-induced T456C alteration in the endogenous mouse *Gars* gene, causing a cysteine-to-arginine switch at residue 201; this produces a range of peripheral nerve defects without affecting survival, reminiscent of CMT2D (12). DRG are heterogeneous collections of neural crest-derived sensory neuron cell bodies found in pairs at each level of the spinal cord, from where they project to and receive information from target peripheral tissues. We chose the initial time point of one month, because the *Gars*^*C201R*/+^ mice are beginning to show overt symptoms, and we have previously performed detailed analyses of their neuromuscular synapses at this age (26).

Thoracic and lumbar DRG sensory neurons were cultured from wild-type and mutant mice, fixed 24 h later, and stained with the pan-neuronal marker *β*III-tubulin to highlight afferent nerve cell somas and processes (Fig. 1A). Mutant cultures showed no difference from wild-type in the percentage of cells bearing neurites (Fig. 1B, top left) or the length of the longest neurite (Fig. 1B, top right); however, there was a significant reduction in the cell body area of *Gars*^*C201R*/+^ neurons (Fig. 1B, bottom left). Cultures were also co-stained with the apoptotic marker activated caspase 3, and average fluorescence intensity per neuron measured at 4, 48, and 96 h post-plating (Fig. 1B, bottom right). There was no difference between genotypes, suggesting that mutant neurons are as healthy as wild-type up to four days in culture, and that cell death *in vitro* is unlikely to be a major contributing factor to the diminished soma area phenotype.

**Figure 1.**
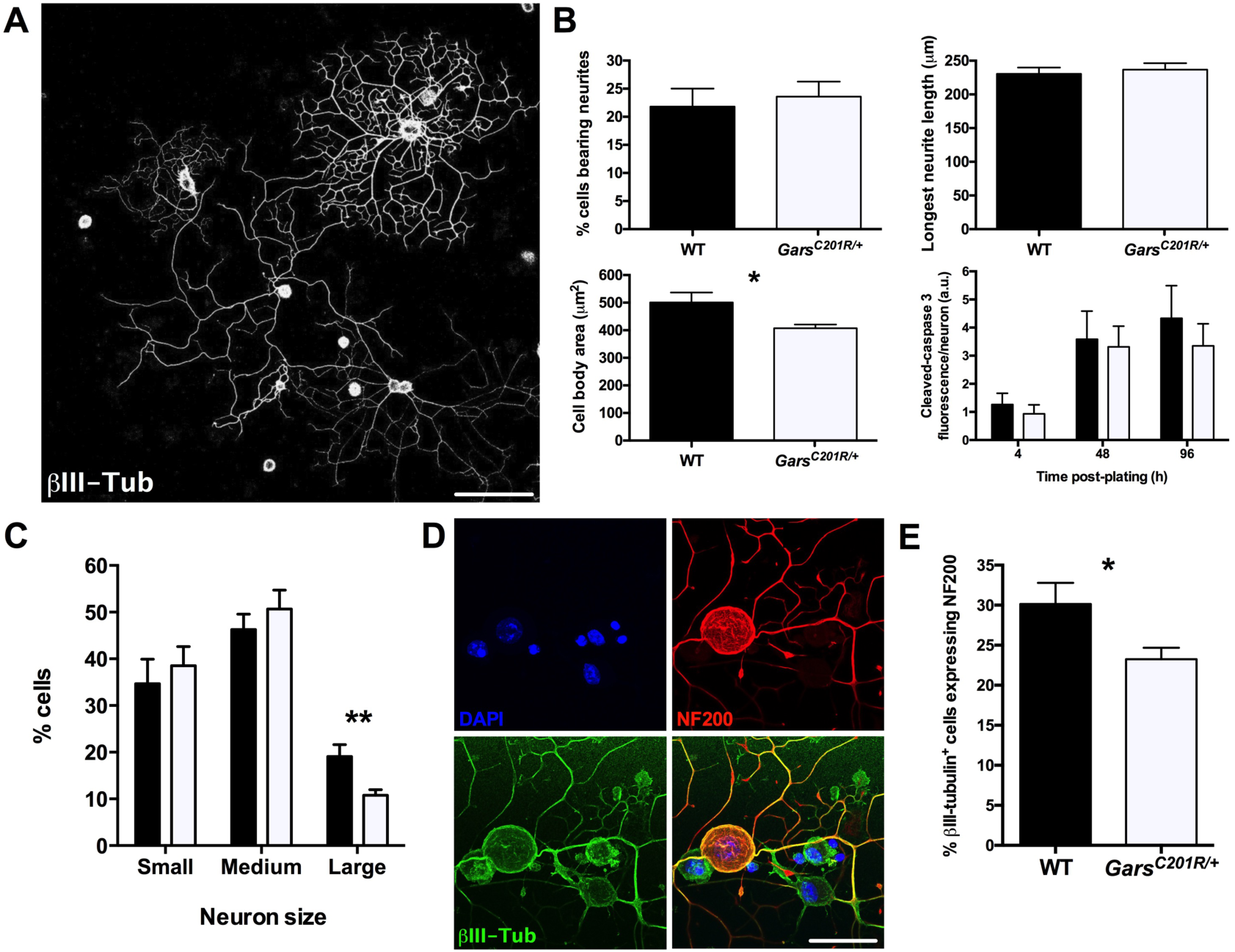
*Gars*^*C201R*/+^ primary DRG cultures have a smaller percentage of large area/NF200^+^ sensory neurons. **(A)** Representative single plane image of one month old wild-type primary dorsal root ganglia (DRG) sensory neurons stained 24 h post-plating with the pan-neuronal marker *β*III-tubulin (white). **(B)** *Gars*^*C201R*/+^ sensory neurons show no difference in the percentage of cells bearing neurites (top left, *P* = 0.678, unpaired *t*-test) or the longest neurite length (top right, *P* = 0.647, unpaired *t*-test), but have a significantly smaller cell body area (bottom left, * *P* = 0.022, unpaired *t*-test). Moreover, mutant cultures do not show signs of cell death above wild-type levels, as assessed by cleaved-caspase 3 staining intensity per neuron (bottom right, two-way ANOVA, *P* = 0.002, time point; *P* = 0.421, genotype; *P* = 0.885, interaction between the two variables). a.u., arbitrary units. **(C)** Mutant DRG cultures possess a significantly lower percentage of large area neurons (cell body area >706 μm^2^, see methods for criteria) than wild-type. ** *P* = 0.008, unpaired *t*-test between % large cells. **(D)** Representative collapsed Z-stack image of DRG neurons stained for DAPI (blue), *β*III-tubulin (green), and the medium-large neuron marker neurofilament 200 (NF200, red). **(E)** Consistent with the reduced percentage of large area neurons (C), *Gars*^*C201R*/+^ cultures have a lower percentage of cells expressing NF200. * *P* = 0.013, Mann-Whitney *U* test. Four (B-C) and six (D) mice/genotype were analysed. Scale bars = 100 μm (A) and 20 μm (E). See also Fig. S1 and 2A.

Sensory neurons can be broadly subdivided into functional classes based on their stimulus response; for example, mechanosensitive neurons that respond to touch, proprioceptive neurons that sense the body’s position in space, and nociceptors that relay noxious stimuli. These classes have been linked to a range of anatomical and physiological characteristics, such as cell soma size, protein markers, and electrophysiological properties, which can be used for reliable functional identification (24, 27). Disparate sensory subtype sensitivities have previously been observed in mouse models of peripheral nerve disease (28, 29). In order to see whether a particular kind of sensory neuron may be preferentially affected by the *Gars C201R* mutation, we divided the *β*III-tubulin^+^ cell bodies into small, medium, and large area neurons based on previously suggested criteria (30). Within these size groups, we again saw no difference between wild-type and mutant neurite length or cell death levels (Fig. S1). However, we did observe a significantly smaller percentage of large area neurons in *Gars*^*C201R*/+^ cultures (Fig. 1C). This result confirms the smaller average mutant cell body area and begins to clarify the etiology of the phenotype, as it could have been due to an increase in small area neurons without large soma neurons being affected.

To differentiate between large and small sensory neurons at the molecular level, and thereby rule out the smaller body size of mutant mice as being the cause of the reduced cell soma area, anti-neurofilament 200 (NF200) was used to mark medium-large neurons with myelinated axons (Fig. S2A), often described as A-fibres (31). Corroborating the cell body measurements, *Gars*^*C201R*/+^ cultures had a significantly smaller percentage of *β*III-tubulin^+^ cells (green) that expressed NF200 (red) than wild-type (Fig. 1D-E). We have thus confirmed at both the morphological and biochemical levels that mutant *Gars* DRG cultures display a significantly reduced percentage of large area neurons.

### Sensory identity is perturbed *in vivo*

To determine whether the *in vitro* sensory phenotypes are also detected *in vivo*, lumbar DRG were dissected from one month old animals, sectioned, and immunohistochemical analysis performed using established markers. Staining for *β*III-tubulin (green, Fig. 2A), we were able to replicate the *in vitro* phenotype of significantly reduced soma size in *Gars*^*C201R*/+^ DRG *in vivo* (Fig. 2B). In addition to NF200, peripherin expression can simultaneously demarcate cell somas of small diameter neurons with thinly myelinated or unmyelinated axons (Aδ- and C-fibres, Fig. S2A) (32), with expression of the two markers being largely mutually exclusive (33). There is some contention as to whether NF200 and peripherin are good indicators of myelination (34); nevertheless, they are well established neuronal size indicators. Anti-NF200 and anti-peripherin were thus used to identify medium-large (red) and small (green) sensory neurons, respectively (Fig. 2C and S2B). *Gars*^*C201R*/+^ DRG show a significantly smaller percentage of NF200-expressing cells (Fig. 2D) and a reciprocal increase in the percentage of peripherin^+^ cells (Fig. 2E). There was only a small degree of co-expression between the two markers (2.3 ± 0.3% versus 2.5 ± 0.4%). The percentage of NF200-expressing wild-type cells is similar to previously reported (35). We have thus shown that the *in vitro Gars*^*C201R*/+^ sensory phenotype of having a smaller percentage of large area/NF200^+^ cells is replicated *in vivo*.

**Figure 2.**
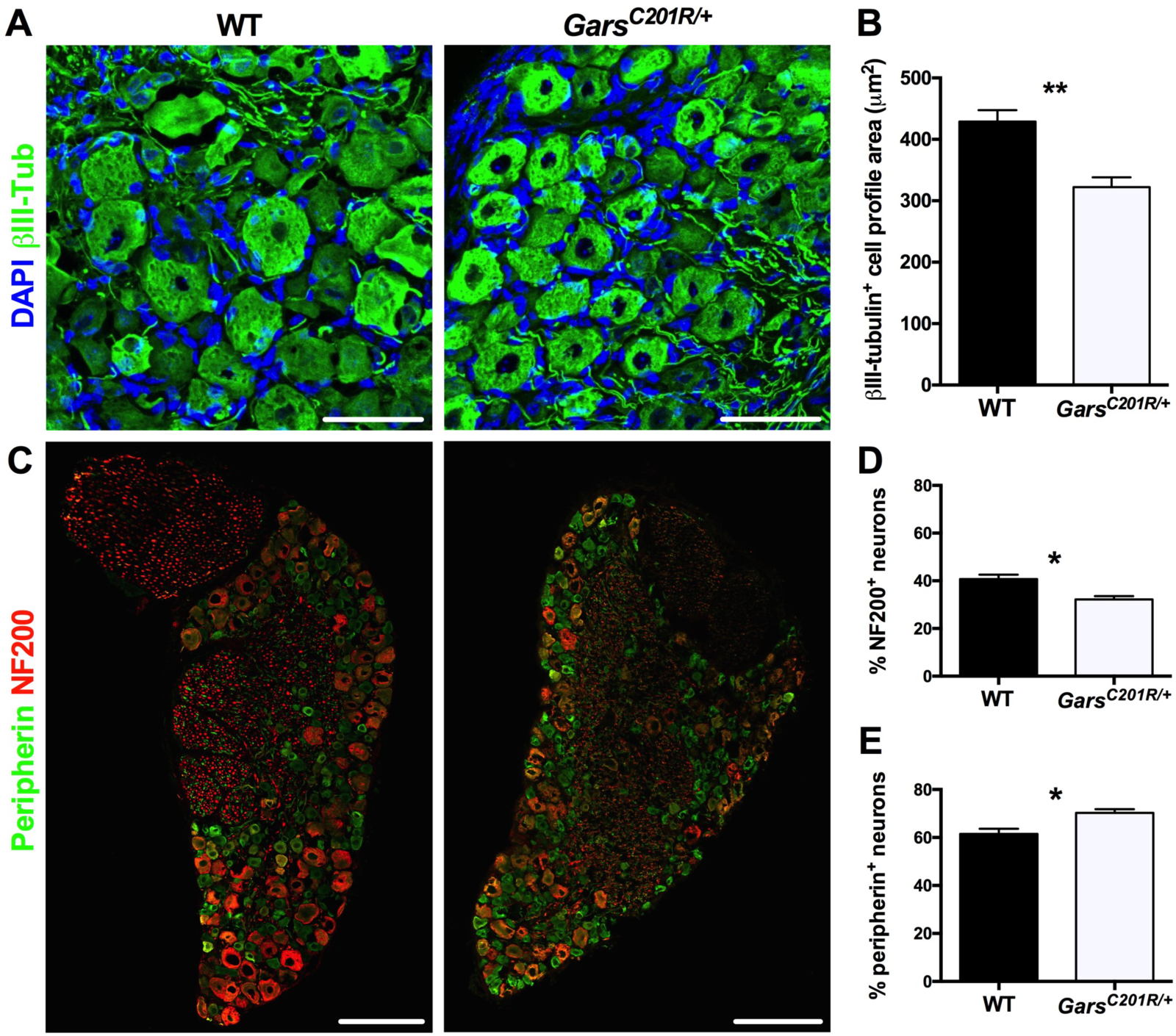
Mutant DRG also have a smaller percentage of large area sensory neurons at one month *in vivo*. **(A)** Representative collapsed Z-stack images of wild-type (left) and *Gars*^*C201R*/+^ (right) DRG at one month stained for DAPI (blue) and the pan-neuronal marker *β*III-tubulin (green). The average cell profile area of mutant sensory neurons is significantly smaller than wild-type. ** *P* = 0.005, unpaired *t*-test. **(C)** Representative wild-type (left) and *Gars*^*C201R*/+^ (right) DRG stained for NF200 (red), marking medium-large sensory neurons, and peripherin (green), labelling small sensory neurons. Images are single confocal planes. **(D-E)** Compared to wild-type, mutant DRG possess a significantly smaller percentage of NF200^+^ cells (D, * *P* = 0.011, unpaired *t*-test) and a concomitant increase in the percentage of peripherin^+^ cells (E, * *P* = 0.015, unpaired *t*-test). Four mice/genotype were analysed. Scale bars = 50 μm (A) and 100 μm (C). See also Fig. S2-5.

To determine whether NF200-expressing cells are selectively affected, DRG sections were tested for the presence of activated caspase 3 (green, Fig. S3A-C). Similar to the *in vitro* results, mutant DRG sections showed no increase in cleaved-caspase 3 signal (Fig. S3B), indicating that differential post-natal cell death is unlikely to be playing a critical role in the reduced percentage of NF200^+^ cells. To test whether mutant ganglia contain increased numbers of peripherin-expressing cells, serial sectioning of L5 DRG was performed (Fig. S3D). The L5 DRG was chosen due to its size and because the resident sensory neurons target distal tissues of the hind limbs, where neuromuscular pathology has been observed in *Gars* mice (11, 15, 26). Counting *β*III-tubulin^+^ (red) cell profiles to estimate the number of neurons per DRG, we found no difference between wild-type and mutant ganglia (Fig. S3E). These profile counts are similar to published approximations from both mouse and rat (36, 37). Given the lack of cell death and similar cell profile counts, the alteration of sensory subtypes in *Gars*^*C201R*/+^ DRG at one month *in vivo* are consistent with a perturbation of neuronal fate.

### The alteration in sensory neuron subtypes correlates with overall disease burden in CMT2D mice

We have previously shown that NMJ pathology correlates with CMT2D severity by comparing *Gars*^*C201R*/+^ with the more severe *Gars*^*Nmf249*/+^ mouse mutant (26, 38), which displays frank denervation, peripheral axon loss, and genetic background-dependent mortality at 6-8 weeks (11). This model has a spontaneous CC-to-AAATA mutation causing proline at residue 278 to be substituted for lysine and tyrosine (11). Similar to the milder allele, one month old *Gars*^*Nmf249*/+^ DRG possessed a significantly lower percentage of NF200^+^ (red) somas (Fig. S4A-B) and a significantly greater percentage of peripherin^+^ (green) neurons compared to wild-type (Fig. S4A and C). When the values from both mutant alleles were compared, *Gars*^*Nmf249*/+^ DRG had a significantly lower percentage of NF200-expressing cells than *Gars*^*C201R*/+^ (Fig. S4B), and a significantly higher percentage of peripherin^+^ cells (Fig. S4C). Importantly, the results hold true when *Gars*^*C201R*/+^ and *Gars*^*Nmf249*/+^ mutant percentage values relative to their respective wild-types are statistically compared for both NF200 (*Gars*^*C201R*/+^, 79.1±3.4% versus *Gars^Nmf249/+^*, 56.6±7.4%) and peripherin staining (*Gars*^*C201R*/+^, 114.2±2.5% versus *Gars*^*Nmf249*/+^, 124.9±4.6%) (data not shown, *P* < 0.05, Sidak’s multiple comparisons test). This indicates that the DRG phenotype correlates with the severity of the *Gars* allele. Moreover, no differences in activated caspase 3 were observed between wild-type and *Gars*^*Nmf249*/+^ ganglia (Fig. S4D), once again suggesting that cell death is unlikely to be a major contributor to this cellular phenotype.

### Mutant mechanoreceptors and proprioceptors are equally affected, as are nociceptor subtypes

Staining for NF200 and peripherin can narrow down sensory neuron classification, but cannot pinpoint function. We therefore used additional markers that broadly relate to the relayed sensory cues. Medium to large area neurons positive for NF200 can be subdivided into two main classes based on the absence or presence of parvalbumin (Fig. S2A). Sensory neurons expressing NF200, but lacking parvalbumin are largely regarded as mechanosensitive cells, whereas those NF200^+^ neurons co-expressing parvalbumin are proprioceptive (24, 25). Parvalbumin also labels a small population of low threshold cutaneous mechanoreceptive neurons, so there is the minor caveat that not all parvalbumin+ neurons are proprioceptive (39). Small area, peripherin-expressing neurons can also be divided into non-peptidergic, principally mechanical nociceptors and peptidergic, mainly thermal nociceptors based on the binding of isolectin B4 (IB4) and the expression of calcitonin gene-related peptide (CGRP), respectively (Fig. S2A) (40-42). However, ablation of CGRP^+^ neurons has an effect on a small proportion of the IB4^+^ population (43). Wild-type and *Gars*^*C201R*/+^ DRG sections were first stained with *β*III-tubulin (blue), NF200 (red), and parvalbumin (green), and the percentage of NF200^+^ cells expressing parvalbumin assessed (Fig. S2C and 5A-B). There was no difference between genotypes in the expression of parvalbumin (Fig. S5C), suggesting that, because there are fewer NF200^+^ cells in mutant DRG, mechanoreceptive and proprioceptive neurons are equally affected by mutant *Gars*. Wild-type and *Gars*^*C201R*/+^ DRG also showed similar percentages of peripherin^+^ (blue) cells either binding IB4 (green) or expressing CGRP (red) (Fig. S5B, D, and S2D), suggesting that different subtypes of nociceptor are also equally affected in mutant mice.

### Peripheral but not central sensory nerve endings are anatomically altered in *Gars*^*C201R*/+^ mice

DRG neurons possess a single axon that projects from the cell body before bifurcating and sending one branch distally to peripheral tissues and another centrally to the dorsal horn of the spinal cord. Given the altered frequencies of large and small area DRG neurons found in CMT2D mice (Fig. 1-2 and S3-5), both distal and central sensory nerve endings were analysed. As mutant ganglia possess fewer NF200^+^ cells, we hypothesised that proprioceptive nerve endings would be impaired. We therefore performed serial transverse sectioning along the entire length of one month old wild-type and *Gars*^*C201R*/+^ soleus muscles, in order to assess muscle spindle number and architecture. Spindles are the highly specialised terminals of proprioceptive neurons important for sensing the state of muscle contraction. Sections were stained with DAPI (blue), SV2/2H3 (green), and laminin (red), to identify nuclei, spindles, and the basement membrane, respectively (Fig. 3A). The SV2/2H3 antibody combination identified spindles, as assessed by their stereotypical architecture, whilst additional antibodies against the classic spindle markers parvalbumin and Vglut1 were ineffective (Table S2). Consistent with the reduced number of NF200^+^/parvalbumin^+^ DRG sensory neurons (Fig. 2 and S5), mutant mice had significantly fewer spindles per soleus muscle (Fig. 3B), while wild-type counts were similar to previously reported (44). Furthermore, we found a dramatic decrease in the percentage of fully innervated spindles (Fig. 3C).

**Figure 3.**
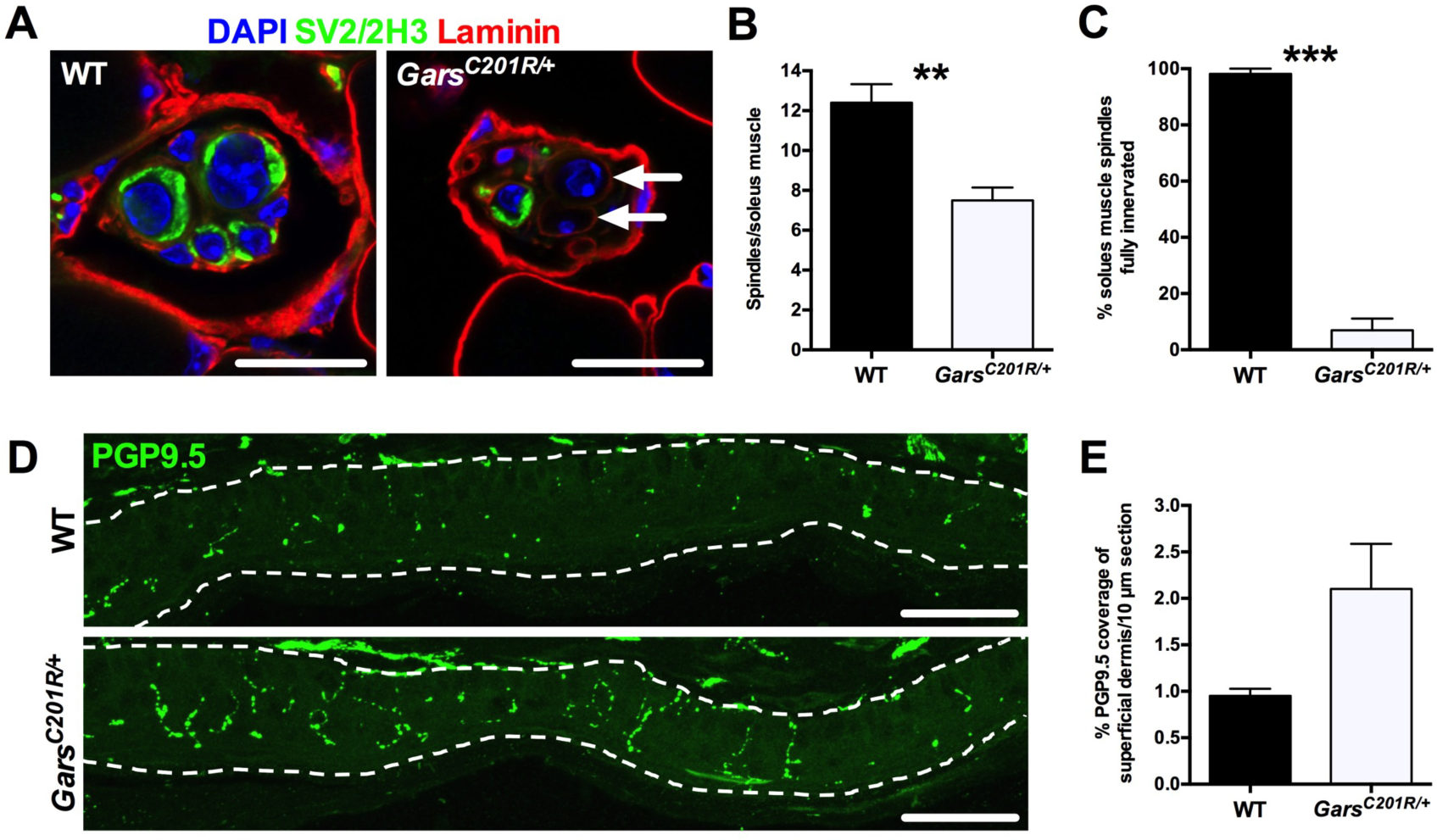
Peripheral nerve endings are altered in *Gars*^*C201R*/+^ mice. **(A)** Representative SV2/2H3^+^ (green) muscle spindles from wild-type (left) and *Gars*^*C201R*/+^ (right) soleus muscles. Anti-laminin highlights the muscle basement membrane (red). N.B., the lack of SV2/3H3 positivity surrounding the central nuclei (DAPI, blue) of the mutant spindle (arrows). Images are single confocal planes. **(B-C)** *Gars*^*C201R*/+^ mice have significantly fewer spindles per soleus muscle (B, ** *P* = 0.005, unpaired *t*-test). Furthermore, mutant spindles display significant denervation (C, *** *P* < 0.001, unpaired *t*-test). **(D)** Representative collapsed Z-stack images taken of the central region of the ventral edge of glabrous hind paw of wild-type (top) and *Gars*^*C201R*/+^ (bottom) mice. Intraepidermal nerve fibres are stained with axonal marker PGP9.5 (green), the epidermis is delineated by dashed lines, and the ventral paw surface is facing down. **(E)** Although not significantly different when tested in isolation (*P* = 0.057, unpaired *t*-test), mutant mice show a significant (*P* < 0.05) increase when multiple time points are included in the analysis and data are tested with Sidak’s multiple comparisons test (see Fig. S8B). 4-5 mice/genotype were analysed. Scale bars = 20 μm (A) and 50 μm (D). See also Fig. S6.

As there are also significantly more peripherin-expressing, pain-sensing neurons in mutant DRG (Fig. 2 and S5), we also assessed nociceptor termini in the skin. Plantar punches of the hind paws were sectioned and stained from one month old mice, and the percentage coverage of the superficial dermis by the axonal marker PGP9.5 assessed (green, Fig. 3D). This method was preferred to intraepidermal nerve fibre counts because it allows a more accurate comparison across different ages. We saw an increase in the peripheral nociceptor innervation in mutant animals (Fig. 3E). Although this did not quite reach significance when tested in isolation, when analysed with data from additional time points, the result was significant (Fig. S8B). The cellular DRG phenotypes of one month old mutant animals therefore correlate with distal proprioceptive and nociceptive sensory neuron deficiencies.

In addition to targeting different peripheral regions for sensing the external environment, sensory neuron subtypes relay their signals to distinct, partially overlapping spinal cord laminae in the dorsal horn. Nociceptors generally form synapses in superficial laminae, numbered I-II, mechanosensitive neurons terminate in deeper laminae III-V, and proprioceptive nerves directly connect centrally and ventrally with interneurons and motor neurons, respectively (25). We therefore sectioned and stained the lumbar spinal cord of one month old mice for the post-synaptic protein PSD95 (green) and the pre-synaptic marker synaptophysin (red) to identify and count synapses in laminae I-III (Fig. S6A-B). Sensory synapses within dorsal laminae IV-V, central, and ventral regions are more widely dispersed and intermingle with a greater number of non-sensory synapses, thus making them more difficult to accurately quantify, so there is the caveat that these analyses do not cover all sensory subtypes. Furthermore, these synapses are not necessarily all sensory. IB4 (blue) was also applied to the sections to aid in the stereotypic anatomical identification of the different laminae. First using PSD95, we saw no difference between wild-type and mutant synaptic density per 100 μm^2^ of lamina I, outer lamina II (IIo), inner lamina II (IIi), or lamina III (Fig. S6C, left). This result was replicated using synaptophysin (Fig. S6C, right), suggesting that despite *Gars* mice having distorted proportions of sensory subtypes in DRG, homeostatic mechanisms regulate afferent entry into the spinal cord in order to maintain consistent synapse numbers.

### Afferent neuron imbalance determines deficits in mutant sensory behaviour

Subtle alterations in the relative abundance of sensory subtypes may or may not cause macroscopic phenotypes and therefore be biologically relevant; we consequently performed four different sensory behavioural tests that broadly depend upon the sensory neuron subtypes that we have assessed in DRG (Fig. S2A). The Von Frey test employs monofilaments of increasing rigidity that are used to apply a specific mechanical stimulus to the hind paws of mice. A response to this test is likely to be mediated, at least in part, by NF200^+^/parvalbumin^−^ neurons. The beam-walking test involves filming mice as they run along a long, thin beam, and then using the videos to assess the percentage of correct foot placements. Amongst other things, this test evaluates the proprioception abilities, and thus the functioning of NF200^+^/parvalbumin^+^ neurons. The Randall-Selitto test assesses a withdrawal response to noxious mechanical stimuli of increasing force either on the hind paw or tail, which requires the activation of mechanical nociceptors, which have been suggested to be non-peptidergic fibres (i.e. peripherin^+^/IB4^+^/CGRP^−^ neurons) (42). Finally, the Hargreaves test examines the function of thermal nociceptors postulated to be the peptidergic fibres (peripherin^+^/IB4^−^/CGRP^+^ neurons) (42), using a noxious heat source on the hind paws and measuring the latency to withdrawal. These four tests were performed on one and three month old wild-type and *Gars*^*C201R*/+^ mice cohorts (Fig. 4 and Tables S3-7). The three month time point was chosen as a later symptomatic age and to provide a useful comparison with previously generated neuromuscular data (26). Concordant with the significantly reduced numbers of NF200-expressing DRG neurons, mutant animals displayed significant defects in reflex withdrawal to a von Frey stimulus at three months and proprioception at both time points (Fig. 4A-B). Moreover, Gars mice showed significant hypersensitivity to both noxious mechanical and thermal stimuli at both one and three months (Fig. 4C-D), consistent with the increased numbers of peripherin^+^ cells in the DRG. When comparing one and three month relative values for *Gars*^*C201R*/+^, only the beam-walking test became progressively worse.

**Figure 4.**
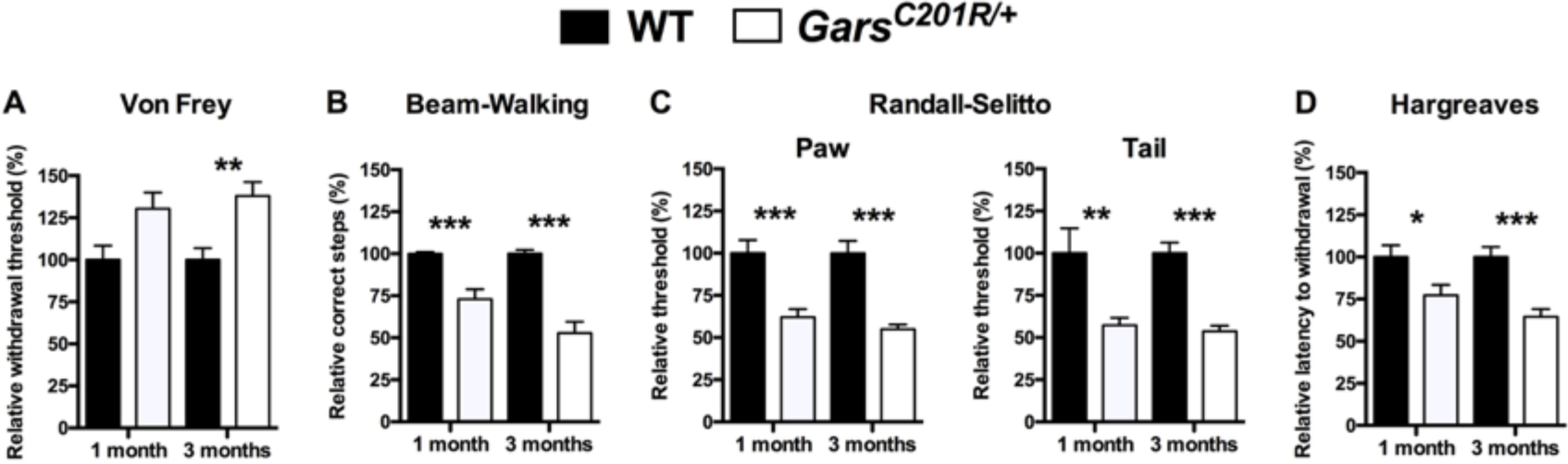
*Gars*^*C201R*/+^ mice display multiple sensory behaviour defects consistent with the distorted DRG cellular phenotype. **(A)** The force required to elicit a response in the Von Frey test is significantly greater for *Gars*^*C201R*/+^ mice, suggestive of a defect in mechanosensation. Two-way ANOVA (*P* < 0.001, age; *P* < 0.001, genotype; *P* = 0.369, interaction). The defect does not worsen over time (*P* = 0.559, unpaired *t*-test). **(B)** In the beam-walking test, mutant mice make significantly more incorrect hind paw steps, perhaps due to defective proprioception. *P* < 0.001, Kruskal-Wallis test, *** P < 0.001 Dunn’s multiple comparison test. This deficiency is exacerbated from one to three months (*P* = 0.030, unpaired *t*-test). **(C)** In stark contrast to the Von Frey test results, mutant mice display hypersensitivity to noxious mechanical stimuli on both the hind paw (*P* = 0.514, age; *P* < 0.001, genotype; *P* = 0.347, interaction, two-way ANOVA) and tail (*P* < 0.001, age; *P* < 0.001, genotype; *P* = 0.138, interaction, two-way ANOVA), as assessed by the Randall-Selitto test. These defects do not worsen with time (*P* = 0.177 and 0.505, unpaired *t*-test). **(D)** Mutant mice also respond faster than wild-type animals to a painful heat source directed to the hind paw, indicative of hypersensitivity to noxious thermal stimuli. Two-way ANOVA (*P* = 0.017, age; *P* < 0.001, genotype; *P* = 0.109, interaction). The defect does not worsen over time (*P* = 0.103, unpaired *t*-test). * *P* < 0.05, ** *P* < 0.01, *** *P* < 0.001, Sidak’s multiple comparisons test (A, C-D). 15 wild-type and 18 *Gars*^*C201R*/+^ mice were analysed in A-B, D, and 11 wild-type and 13 *Gars*^*C201R*/+^ mice were analysed in C. The statistical tests represented on the figures were performed on raw data (Tables S3-7), while the percentages relative to wild-type, which are plotted, were used to compare mutant progression over time. See also Fig. S7 and Tables S8-9.

We also performed motor behaviour testing at the same time points, in order to see whether motor deficits may be contributing to the observed sensory behaviour phenotypes (Fig. S7 and Tables S8-9). Grip strength tests were performed to assess fore and hind limb muscle force and the accelerating Rota-Rod was implemented to measure the complex relationship between motor ability, balance, coordination, and proprioception. We found that both female and male mutant mice showed significant defects in both tests, but, like the sensory behaviours, these did not appear to worsen with age. These motor test results suggest that motor deficiencies may indeed contribute to the mechanosensation and proprioception deficits seen in the *Gars*^*C201R*/+^ mice (Fig. 4A-B). However, given that the beam-walking deficit, but not the grip strength defect, is progressive from one to three months, it appears as though the defective proprioception is partially independent of motor impairment. Furthermore, given that mutant animals are responding quicker to noxious stimuli (Fig. 4C-D), the motor defects are unlikely to be integral to the pain hypersensitivity. It is worth emphasising that the mutants showed a previously unreported phenotype of reduced mechanosensation (Fig. 4A) with the contrasting enhancement of mechanical nociception (Fig. 4C). In summary, the behavioural testing shows that *Gars* mice display multiple disturbances of sensory behaviour that correlate with the cellular phenotypes observed in DRG.

### *Gars*^*C201R*/+^ mice display developmental sensory deficits

In order to see whether the cellular sensory phenotype gets progressively worse with time, we analysed DRG from one day (postnatal day 1, P1) and three month old mice. We were again able to demonstrate at both time points the presence of significantly fewer mutant NF200^+^ neurons (Fig. 5A) and more peripherin^+^ cells (Fig. 5B), confirming the result at one month. Comparing the percentages of NF200^+^ and peripherin^+^ cells in mutant samples relative to wild-type, we see no significant differences between any of the time points (*P* > 0.05, Sidak’s multiple comparisons test). We have thus shown that the disturbed population of sensory neuron subtypes resident in the mutant DRG are present at birth and do not change by early adulthood.

**Figure 5.**
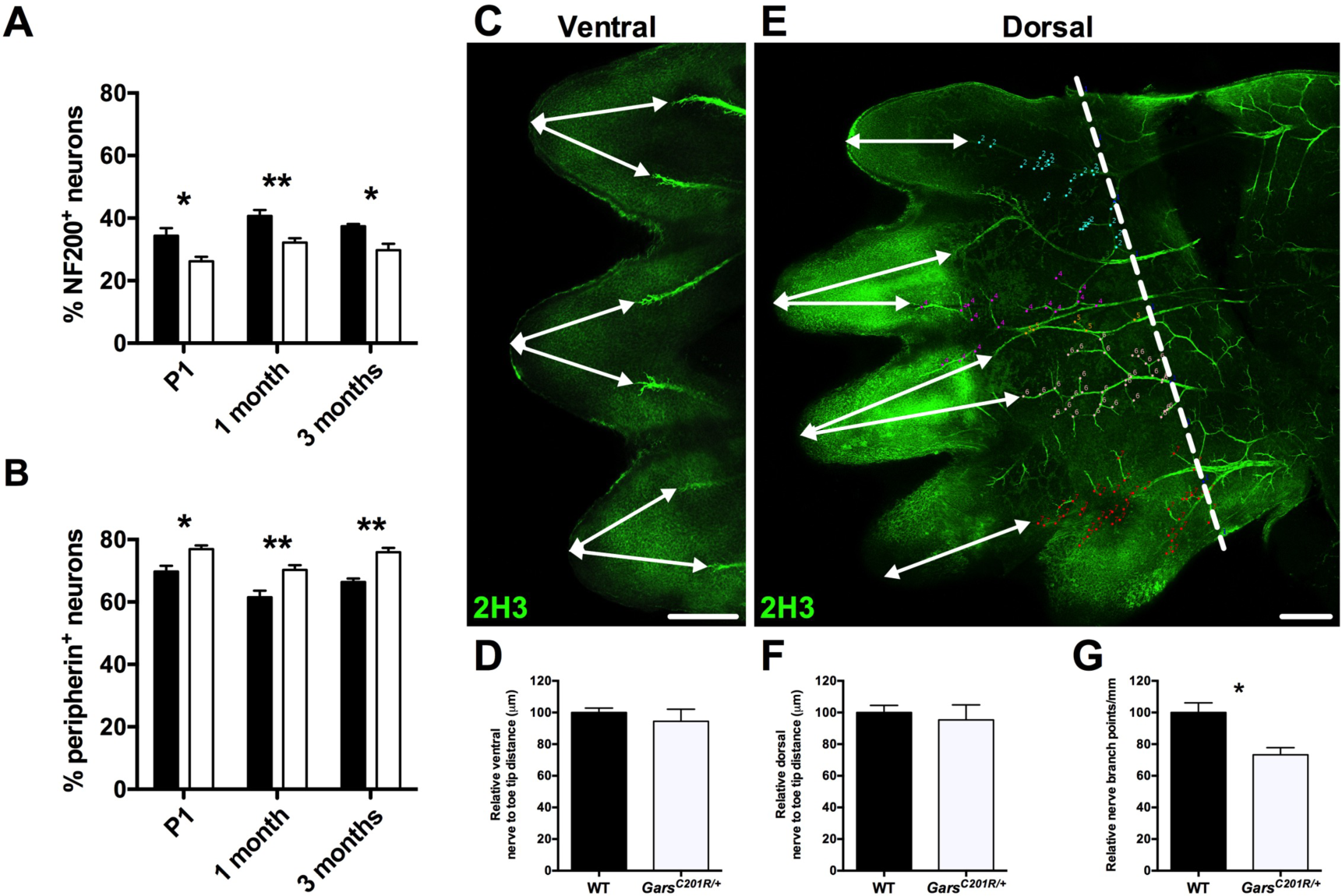
*Gars*^*C201R*/+^ sensory neurons display developmental defects. **(A)** *Gars* mutant DRG display significantly smaller percentages of NF200^+^ cells at P1, one month, and three months, suggestive of a non-progressive, pre-natal defect. The average percentage of NF200^+^ cells in mutant DRG is 74.4% (P1), 79.1% (one month), and 79.4% (three months) relative to wild-type. Two-way ANOVA (*P* = 0.011, age; *P* < 0.001, genotype; *P* = 0.973, interaction). **(B)** Mutant DRG show a reciprocal increase in the percentage of cells expressing peripherin at all three time points. The mean percentage of peripherin^+^ cells in mutant DRG relative to wild-type is 110.9% (P1), 114.2% (one month), and 114.3% (three months). Two-way ANOVA (*P* < 0.001, age; *P* < 0.001, genotype; *P* = 0.769, interaction). Statistical analyses are performed on raw data and not percentages relative to wild-type (A-B). **(C, E)** Representative single confocal plane, tile scan images of ventral (C) and dorsal (E) aspects of an E13.5 hind paw stained for neurofilament (2H3, green). The arrows depict distances from the major nerve branches to the tips of the toes measured in D and F, and the dashed line the dorsal foot plate in which branching density was assessed in G. Scale bars = 250 μm. **(D, F-G)** There was no difference between wild-type and mutant mice in the targeting of sensory nerves to the hind paw extremities on ventral (D, *P* = 0.413, unpaired *t*-test) or dorsal sides (F, *P* = 0.629, unpaired *t*-test). However, *Gars*^*C201R*/+^ neurons display reduced branching in the dorsal foot plate (G, * *P* = 0.0376, unpaired *t*-test). Statistical analyses were performed on percentage values relative to the wild-type mean. 3-9 mice/genotype per time point were analysed. See also Fig. S8.

Cleaved-caspase 3 levels also did not differ, suggesting that cell death is playing no major role in the onset and/or maintenance of this phenotype (Fig. S8A). We also assessed intraepidermal nerve fibre density at P1 and three months. Contrasting with the one month data, we saw no difference between wild-type and mutant at these early and late time points (Fig. S8B). Innervation density declines over time in both mutant and wild-type animals; however, it appears to take longer in the *Gars*^*C201R*/+^ mice.

In order to confirm whether sensory nerve development is affected in *Gars* mutant mice, we analysed axonal projections of small diameter sensory neurons in wholemount hind paws of E13.5 embryos (45, 46). To assess axonal extension, we measured the distance from the main nerve trunk termini innervating the foot plate to the tips of the embryonic digits (Fig. 5C, E). We saw no difference in either the ventral (Fig. 5D) or the dorsal (Fig. 5F) nerve, indicating that nerve terminal extension is unaffected. However, we found that mutant nerves display a significant reduction in branch density in the dorsal floor plate (Fig. 5G). This suggests that arborisation of mutant TrkA^+^ sensory neurons is impaired, and that *Gars*^*C201R*/+^ mice display developmental perturbations in the sensory nervous system.

### Mutant thermal nociceptors display greater excitability

Cell autonomous differences in neuronal excitability (12) may contribute to the pain hypersensitivity phenotype of *Gars* mice (Fig. 4). We therefore cultured DRG neurons from one month old animals and performed calcium imaging experiments using the ratiometric calcium indicator fura-2 (47). We saw no difference in the baseline fura-2 ratio between wild-type and *Gars*^*C201R*/+^ sensory neurons (data not shown, 0.840 ± 0.012 versus 0.835 ± 0.014, *P* = 0.787, unpaired *t*-test), suggestive of equivalent resting state calcium levels in wild-type and mutant neurons. When 50 mM KCl was applied to the cells to trigger depolarisation, there was also no difference in the elicited response (Fig. 6A). In these live DRG cultures, NF200^+^ and peripherin^+^ neurons cannot be readily differentiated. We therefore applied 1 μM capsaicin, which activates the non-selective cation channel TRPV1 (48), in order to functionally differentiate thermal nociceptors. Addition of capsaicin precipitated a greater relative change in the fura-2 ratio of capsaicin-responsive *Gars*^*C201R*/+^ than wild-type neurons (Fig. 6B), and this appears to be a general defect rather than a larger response from a sub-selection of capsaicin-responsive neurons. These experiments therefore indicate that mutant thermal nociceptors are intrinsically hyper-responsive to painful stimuli, which is likely to contribute to the pain hypersensitivity phenotype observed in adult *Gars*^*C201R*/+^ mice.

**Figure 6.**
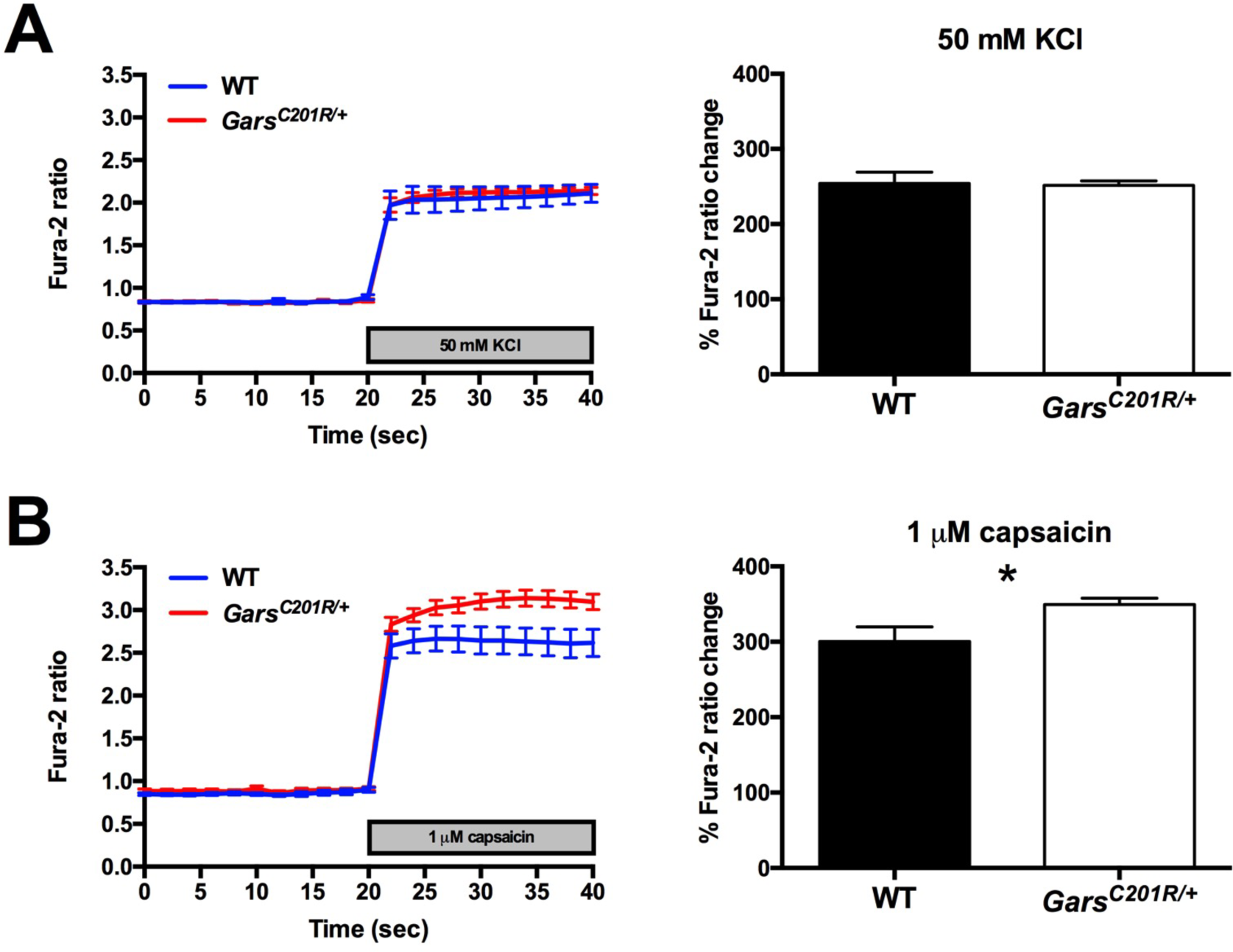
Thermal nociceptors from mutant *Gars* mice are hyperexcitable. **(A)** One month old wild-type (blue) and mutant (red) primary DRG neurons show no difference in their responses to 50 mM KCl 24 h post-plating (right, *P* = 0.864, unpaired *t*-test), as assessed using the ratiometric calcium indicator fura-2. **(B)** The increase in cytosolic calcium concentration upon stimulation by 1 μM capsaicin is greater in *Gars*^*C201R*/+^ than wild-type neurons. Only data generated from capsaicin-responsive cells (i.e. thermal nociceptors) are included in these graphs. Wild-type and mutant cells display similar baseline calcium levels, but capsaicin triggers a significantly larger increase in the fura-2 ratio (340 nm:380 nm) in *Gars*^*C201R*/+^ neurons (right, * *P* = 0.0236, unpaired *t*-test).

## Discussion

CMT2D patients display both motor and sensory pathology, yet the sensory component has received little attention both in humans and animal models. We therefore performed a detailed examination of the sensory system of CMT2D mice in order to better understand the afferent nerve pathogenesis (see Fig. 7 for an overview). We found that mutant DRG possess fewer large diameter, NF200^+^ cells and a concomitant increase in the number of small diameter, peripherin^+^ neurons (Fig. 1-2), a phenotype that nicely correlates with CMT2D mutant severity (Fig. S4), and alterations in sensory behaviour (Fig. 4). Assessment of activated caspase 3 levels and DRG neuron counts indicate that this phenotype is unlikely to be caused by post-natal cell death or defective neural crest migration and survival, but is rather a developmental sensory subtype switch (Fig. S3). Consistent with pre-natal onset, the DRG phenotype is present at birth (Fig. 5A-B). Moreover, embryonic sensory nerve branching in the mutant hind paw is defective (Fig. 5G), similar to the previously reported impairment in facial motor neuron migration (18). Using several markers for sensory function, we observed that mechanoreceptive and proprioceptive neurons are equally affected by *Gars* mutation, as are non-peptidergic and peptidergic nociceptors (Fig. S5). The pathological effect of mutant GlyRS could therefore be triggered by the differential expression of specific genes integral to sensory diversification between the mutually exclusive NF200^+^ and peripherin^+^ neuronal populations (e.g. tropomyosin receptor kinase (Trk) receptors) (49). Differences in cellular origin or timing of gene expression leading to subtype specification could also contribute to the DRG phenotype (50, 51).

**Figure 7.**
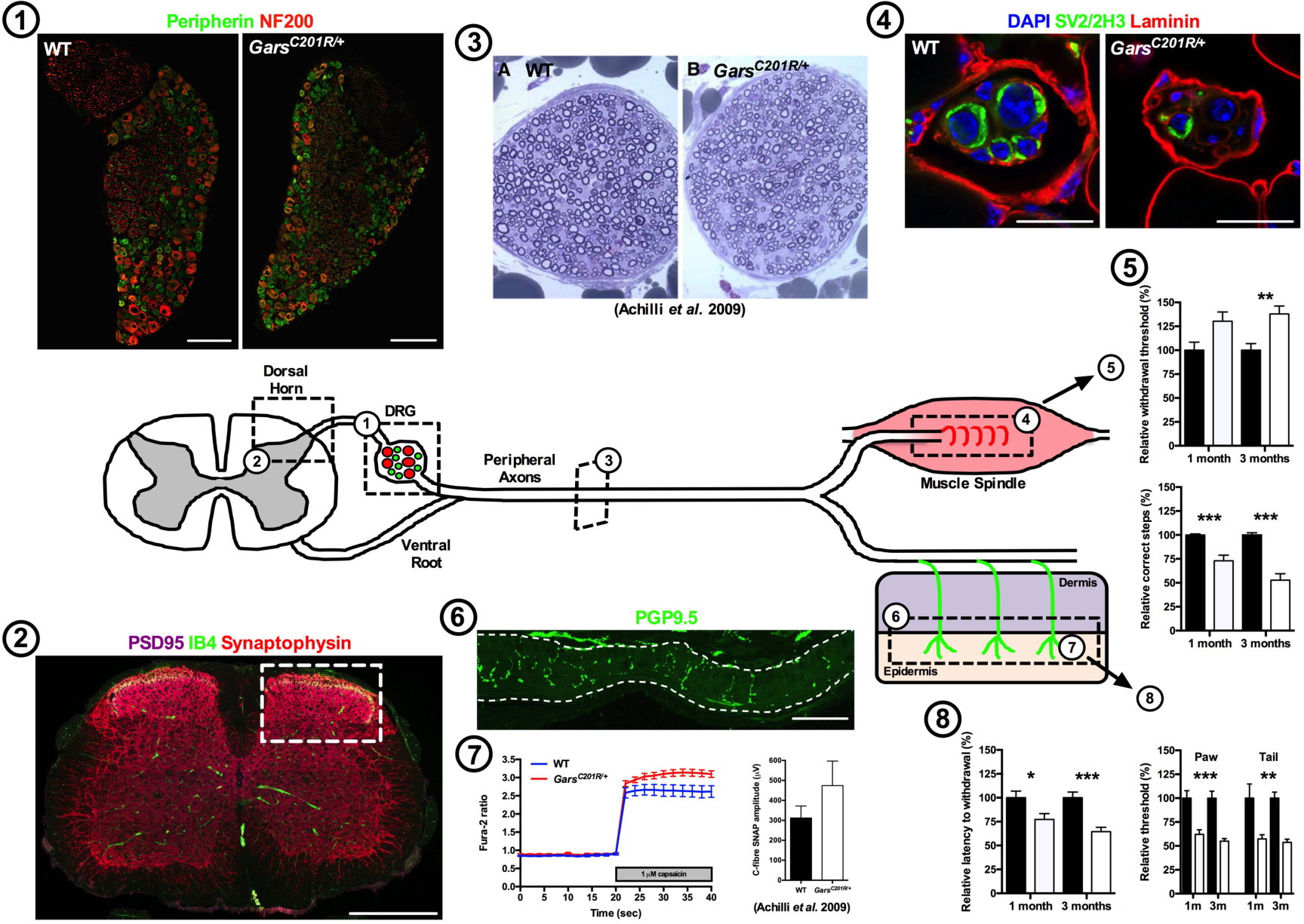
CMT2D sensory nervous system pathology. *Gars* mutations distort the proportions of sensory neuron subpopulations in the DRG, such that mutant mice have fewer medium-large (red, NF200^+^) and more small (green, peripherin^+^) area neurons from birth to at least three months (1). This does not affect synapse numbers within distinct spinal cord dorsal laminae (2), but may account for the previously reported (12) reduced axon calibre of sensory nerves (3) and the diminished number and innervation of muscle spindles in the soleus (4), both of which likely contribute to mutant mechanosensation and proprioception deficiencies (5). Furthermore, mutant *Gars* nociceptors show delayed pruning in peripheral tissue (6), hyperexcitability (7, left), and the previously reported (12) increased sensory nerve action potential (SNAP) amplitude (7, right). These phenotypes may be pertinent to the pain hypersensitivity observed in *Gars* mice (8). Together, our work indicates that CMT2D may arise from the intricate interaction between subverted neurodevelopment and neurodegeneration. Scale bars = 100 μm (1), 50 μm (2 and 6), and 20 μm (4). The unaltered pictures of transverse sensory nerves (3) and graphically-depicted SNAP data (7) were taken from Achilli *et al*. 2009 with permission under the Creative Commons Attribution (CC-BY 3.0) license (http://creativecommons.org/licenses/by/3.0/).

CMT2D-associated mutant GlyRS was recently shown to aberrantly bind to and antagonise the neuronal receptor protein NRP1 (18). Although NRP1 was the focus of that study, mutant GlyRS was shown to aberrantly interact with a number of other proteins found on the neuronal surface, albeit to a lesser degree (18). One of these proteins was TrkB, a neurotrophin receptor that, once activated, specifically drives differentiation and survival of mechanosensitive sensory neurons (52). Similarly, TrkA and TrkC are pivotal to the survival of nociceptive and proprioceptive nerves, respectively (53, 54). A previous study has shown that expression of TrkC from the *TrkA* locus caused a developmental fate switch in DRG sensory subtypes (55). Given that GlyRS is expressed during early development (11, 12), and that arborisation of TrkA^+^ neurons is developmentally impaired in CMT2D mice (Fig. 5G), it is not inconceivable that a low level of mutant GlyRS binding to TrkB, and possibly other Trk receptors, could subtly affect their ability to signal or induce transactivation, and thus subvert sensory neuron differentiation and/or survival during early stages of development. Interestingly, *Gars*^*C201R/C201R*^ homozygous mutants display increased DRG cell numbers compared to littermate controls (12), which may indicate that a higher dose of mutant protein could have a greater effect on the sensory system

Regardless of the cause of the afferent imbalance in mutant DRG, it is clear that it represents a major, non-progressive, developmental component of the sensory phenotype of CMT2D mice. This is in agreement with the sensory phenotype not worsening from one to three months (except for proprioception, Fig. 4), and consistent with *Gars*^*C201R*/+^ sensory saphenous nerve showing a smaller average axon caliber, but no signs of degeneration or axon loss up to three months (12). This non-progressive sensory insult might also be seen in humans, as the extent of CMT2D patient sensory deficiency is reported to be reliant not upon disease duration, but severity (19). Accordingly, without the initial developmental perturbation of the sensory system, afferent pathology may not arise, which could explain the predominantly motor presentation of dSMA-V patients. An element of mutant *GARS*-related sensory pathology may therefore be binary; if mutant GlyRS triggers the initial developmental insult, CMT2D will arise, but if not, then dSMA-V manifests.

In addition to a prenatal developmental disturbance, maturation and degenerative pathways are also contributing to GlyRS-mediated pathology. *Gars*^*C201R*/+^ mice possess significantly fewer muscle spindles and reduced innervation per spindle (Fig. 3B-C), which is probably reflective of reduced formation during development and subsequent degeneration. Together with the previously reported decrease in amplitude of sensory nerve action potentials (SNAPs) in large area neurons (1.7±0.2 μV versus 1.2±0.2 μV) (12), both defects are likely to contribute to the defective proprioception, while progressive distal nerve deterioration perhaps accounts for proprioception being the only sensory behaviour to decline over time (Fig. 4B). Therefore, it is conceivably not a coincidence that the ability of CMT2D patients to sense vibration is the most impaired sensory symptom.

We have previously shown that a developmental delay in NMJ maturation precedes synaptic degeneration in *Gars* mouse distal muscles (26). Interestingly, we see a similar pruning deficiency in the intraepidermal nociceptors of the mutant hind paws (Fig. S8B). We believe that this represents impairment of the early post-natal refinement of sensory architecture (56) (akin to the motor phenotype) as opposed to degeneration, as the latter would likely precipitate a reduction in the pain hypersensitivity phenotype by three months. To find an alternate explanation, we performed synapse counts in distinct spinal cord dorsal laminae (Fig. S6) and calcium imaging experiments on primary DRG cultures (Fig. 6). We saw no difference between genotypes in dorsal horn synapse densities (Fig. S6C). This suggests that homeostatic mechanisms are at work to restrict C-fibre entry into the spinal cord and that there is perhaps an excess of NF200^+^ neuronal branches targeting dorsal laminae in wild-type mice. Nevertheless, dorsal horn synapse counts do not assess synaptic strength and therefore it is uncertain whether or not central sensitisation has occurred. To assess this peripherally, we analysed cytosolic calcium dynamics, and found that mutant thermal nociceptors are more responsive to capsaicin than wild-type neurons (Fig. 6B). The increased number of small area neurons and axons probably account for the previously reported (non-significant) increase in mutant C-fibre SNAP amplitude (312±60 μV versus 474±123 μV) (12). Through activity-dependent mechanisms of peripheral or central plasticity, such as differential ion channel expression/phosphorylation or synaptic potentiation (57), we hypothesise that this, in turn, could alter neuronal excitability and at least partly explain the inherent thermal nociceptor hyperexcitability and the pain hypersensitivity phenotypes.

In summary, we have shown that CMT2D mice display numerous sensory symptoms that hinge upon a disturbed equilibrium between functional subtypes of afferent neurons. This phenotype is likely developmental in origin and could serve to explain the variable sensory pathology of *GARS*-associated neuropathy. In light of the range of deficits reported in *Gars* mice both here and elsewhere, we propose that CMT2D pathology reflects a complex interplay between developmental, maturation, and survival pathways, a conclusion that has profound implications for the development of novel therapies and timing of therapeutic intervention for the treatment of this disease.

## Materials and Methods

### Animals

*Gars*^*C201R*/+^ and *Gars*^*Nmf249*/+^ mice were maintained as heterozygote breeding pairs on a predominantly C57BL/6 background as described previously (11, 12, 58). Mice sacrificed for the one month and three month time points were 28-36 and 88-97 days old, respectively. Post-natal day 1 (P1) was designated as the day following the day a litter was first found. To reduce the overall number of mice used, behavioural testing was performed on and multiple tissues were harvested from both males and females for this and other parallel studies (59). *Gars*^*C201R*/+^ handling and experiments were performed under license from the United Kingdom Home Office in accordance with the Animals (Scientific Procedures) Act (1986), and approved by the University College London – Institute of Neurology Ethics Committee for work in London and by the University of Oxford Ethical Review Panel for experiments conducted in Oxford. *Gars*^*Nmf249*/+^ mouse husbandry and procedures were conducted in accordance with the NIH Guide for Care and Use of Laboratory Animals and approved by The Jackson Laboratory Animal Care and Use Committee. Ear clips were used to extract DNA as previously described (60), and animals were genotyped using previously published protocols (11, 12, 58).

### DRG dissection and culturing

Ethanol-sterilised 12 mm coverslips (VWR International, Radnor, PA, MENZCB00120RAC20) placed in 24-well plates (Corning, New York, NY, 3524) were treated with 20 μg/ml poly-D-lysine (Becton Dickinson, Franklin Lakes, NJ, 354210) in Ca^2+^/Mg^2+^-free Hank’s Balanced Salt Solution (HBSS, Life Technologies, Carlsbad, CA, 14170) for at least 12 h at 4°C. Wells were washed three times with Ca^2+^/Mg^2+^-free Dulbecco’s phosphate buffered saline (DPBS, Life Technologies, 14190), and thoroughly dried. 10-20 μl HBSS containing 20 μg/ml laminin (Sigma Aldrich, St. Louis, MO, L2020) was pipetted onto the centre of coverslips to concentrate the neurons and incubated at 37°C for 4-6 h. DPBS was pipetted between wells to restrict evaporation of the laminin solution. DRG neurons used in calcium imaging experiments were plated in 8 well μ-slides (Ibidi, Martinsried, Germany, 80826) treated as above without coverslips. Before starting the dissection, a previously prepared and frozen collagenase/dispase enzyme solution was thawed at 37°C for no longer than 2 h before use; 24 mg collagenase type II (1 U/μl, Worthington Biochemical Corporation, Lakewood, NJ, 4176 or Life Technologies, 17101015) and 28 mg dispase II (Sigma, D4693) were added to 6 ml Ca^2+^/Mg^2+^-free HBSS, and filter-sterilised through a 0.22 μm filter (Appleton Woods, Birmingham, UK, FC121) before aliquotting and freezing. DRG were dissected as previously described (61). To limit technical variability, 20-24 DRG per animal were dissected from thoracic (T1-T13) and lumbar (L1-L5) spinal cord regions of one wild-type and one *Gars*^*C201R*/+^ mouse during the same culturing session. DRG were enzymatically digested at 37°C for 10 min in collagenase/dispase, before manual dissociation in cell media using a series of fire-polished glass Pasteur pipettes (VWR, 612-1701) of descending bore size. Cells were finally spun down at 1,000 × *g* for 5 min, before resuspension. The laminin solution was pipetted off the coverslips/μ-slides and a small volume of cells immediately added. Wild-type and mutant cells were plated at similar densities. Cells were kept at 37°C in a 5% (v/v) CO_2_ humidified atmosphere. After 1-2 h, plate and μ-slide wells were carefully flooded with media to a total volume of 500 μl and 200 μl, respectively. F12 media + L-Glutamine (Life Technologies, 11765) was supplemented with 10% (v/v) foetal bovine serum (Life Technologies, 10270), 1% (v/v) penicillin-streptomycin (10,000 U/mL, Life Technologies, 15140), and freshly added 20 ng/ml mouse glial cell line-derived neurotrophic factor (Peprotech, Rocky Hill, NJ, 450-44) with 2 mg/ml bovine serum albumin (BSA, Sigma, 10735094001) carrier protein in water. For calcium imaging experiments, cultures were also supplemented with 50 ng/ml mouse nerve growth factor *β*(Peprotech, 450-34) with 2 mg/ml BSA in water.

### Cell and tissue immunohistochemistry

All steps were completed at room temperature, apart from overnight incubations, which were conducted at 4°C. For cell immunohistochemistry, DRG media was carefully aspirated 24 ± 1 h post-plating, and cells fixed with 4% (w/v) paraformaldehyde (PFA, Electron Microscopy Sciences, Hatfield, PA) for 20 min; 16% PFA stock was diluted in phosphate buffered saline (PBS, 137 mM NaCl [Sigma, S3014], 10 mM Na_2_HPO_4_ [Sigma, S3264], 2.7 mM KCl [Sigma, P9541], 1.8 mM KH_2_PO_4_ [Sigma, P9791]) to achieve the final working solution. Neurons were permeabilised for 30 min using 0.3% (w/v) Triton X-100 in PBS, before blocking for 30 min in permeabilisation solution containing 5% (w/v) BSA, and probing overnight with primary antibodies (see below) in block solution. The following day, coverslips were washed three times for 10 min in PBS, probed with secondary antibodies (see below) and 4’,6-diamidino-2-phenylindole, dihydrochloride (DAPI, Life Technologies, D1306) for 2 h, and washed three times with PBS, before mounting on slides in fluorescence mounting medium (Dako, Glostrup Municipality, Denmark, S3023). Slides were kept in the fridge and allowed to set overnight before imaging.

For tissue immunohistochemistry, L1-L5 DRG were dissected and fixed in 4% PFA for 2 h, before washing in PBS, and equilibrating in 20% (w/v) sucrose (Sigma, S7903) in PBS overnight. Plantar punches were collected from the right hind paw using a 5 mm punch (Sigma, Z708771), fixed in 4% PFA overnight, placed in decalcifying solution (15% sucrose, 10% [w/v] ethylenediaminetetraacetic acid [EDTA, Sigma, E5134], 0.07% [w/v] glycerol [Sigma, G5516] in PBS) overnight, and treated with 20% sucrose in PBS overnight (62). Soleus muscles and spinal cords were dissected from mice transcardially perfused with 4% PFA at a rate of ≈3 ml/min for 4-5 min. Soleus muscles were post-fixed for 2 h, before washing with PBS and leaving in 20% sucrose overnight (44). Spinal cords were post-fixed overnight, washed with PBS, and placed in 20% sucrose overnight. The embryonic day 18.5 (E18.5) *Kidins220^−/−^* brain, used as a positive control for activated caspase 3 staining (63), was dissected in PBS, fixed for 2 h in 4% PFA, washed in PBS, and equilibrated in 20% sucrose in PBS overnight. All sucrose-treated tissues were embedded in Tissue-Tek O.C.T. Compound (Sakura Finetek, Torrance, CA, 4583), frozen on dry ice-chilled methanol (Sigma, 32213), and kept at −80°C. 10 μm L1-L5 DRG, one and three month plantar punch, and E18.5 brain sections, 12 μm soleus muscle sections, 20 μm P1 plantar punch sections, and 30 μm spinal cord sections were cut with an OTF Cryostat (Bright Instruments, Huntingdon, UK) and collected on polysine-coated slides (VWR, 631-0107). Transverse sections were cut from all tissues, except for the E18.5 *Kidins220^−/−^* brain (coronal), and the DRG, which were sectioned in stochastic orientations due to their spherical nature. Spinal cords were sectioned onto 12 parallel slides, L1-L5 DRG and the E18.5 brain onto eight parallel slides, soleus muscles and plantar punches on to four parallel slides, and all L5 DRG sections for neuron cell counts were collected onto 1-2 slides/DRG. All sections were air dried for 30-60 min before staining or freezing at −80°C, except for spinal cord sections, which were air dried overnight.

For tissue staining, sections were encircled with a hydrophobic barrier pen (Dako, S2002) on microscope slides, permeabilised three times with 0.3% Triton X-100 in PBS for 10 min, and blocked for 1 h in 10% BSA and 0.3% Triton X-100 in PBS. To facilitate labelling of synaptic structures, spinal cord sections were incubated for 30 min in 50% (v/v) ethanol in water, followed by 10 min in hot (>95°C) citrate-EDTA buffer (10 mM citric acid [Sigma, W302600], 2 mM EDTA [Sigma, ED2SS], pH to 6.2), 10 min in 4 μg/ml Proteinase K (Merck Millipore, Billerica, MA, 124568) in PBS containing 0.3% Triton X-100, and 10 min in 1 mg/ml pepsin (Dako, S3002) in 0.2 M HCl (Sigma, 258148) at 37°C. Samples were then probed overnight with primary antibodies (see below). The following day, sections were processed in the same way as primary DRG neuron coverslips (see above), bathed in fluorescence mounting medium, and covered with 22 × 50 mm cover glass (VWR, 631-0137).

E13.5 hind feet were removed from embryos between the ankle and knee joints and processed, with subtle modifications, as previously described (45). Briefly, feet were fixed for 4 h in 4% PFA, followed by overnight bleaching in 15% hydrogen peroxide (Sigma, H1009) and 1.5% dimethyl sulphoxide (DMSO, Sigma, D1435) in methanol. The following day, feet were moved to 10% DMSO in methanol overnight, before application of primary antibody (see below) in block solution (5% normal goat serum [Sigma, G9023] and 20% DMSO in PBS) for five days. Feet were subsequently washed five times in PBS for 1 h, before applying secondary antibody in block overnight. Finally, feet were washed three times in PBS for 1 h before mounting on microscope slides in Dako medium, and covering with 22 × 50 mm cover glass.

The following primary antibodies were used (Tables S1-2): sheep anti-CGRP (1/200, Enzo Life Sciences, Farmingdale, NY, BML-CA1137), rabbit anti-cleaved-caspase 3 (1/500, Cell Signalling Technology, Danvers, MA, 9661), rabbit anti-laminin (1/1000, Sigma, L9393), mouse anti-neurofilament (2H3, 1/50 or 1/250, developed by Thomas M. Jessell and Jane Dodd, Developmental Studies Hybridoma Bank, Iowa City, IA, supernatant), mouse anti-NF200 (1/500, Sigma, N0142), rabbit anti-parvalbumin (1/1000, Swant, Marly, Switzerland, PV27), rabbit anti-peripherin (1/500, Merck Millipore, AB1530), rabbit anti-PGP9.5 (1/1000, UltraClone, Isle of White, UK, 31A3), rabbit anti-PSD95 (1/200, Frontier Institute, Ishikari, Japan, Af628), mouse pan anti-synaptic vesicle 2 (SV2, 1/250, developed by Kathleen M. Buckley, Developmental Studies Hybridoma Bank, Iowa City, IA, concentrate), guinea pig anti-synaptophysin (1/200, Frontier Institute, Af300), chicken anti-*β*III-tubulin (1/500, Abcam, Cambridge, UK, ab41489), and mouse anti-*β*III-tubulin (1/500, Covance, Princeton, NJ, mms-435P). Tissues were also sometimes incubated with 1 mg/ml isolectin B_4_ (IB4) biotin conjugate from *Bandeiraea simplicifolia* (Sigma, L2140) in PBS at 1/250. The following day, combinations of AlexaFluor secondary antibodies (Life Technologies, A-11001, A-11008, A-11034, A-11039, A-11074, A-21235, A-21236, A-21245, A-21424, A-21429, A-21436) at 1/1000, 2 mg/ml streptavidin, Pacific Blue conjugate (Life Technologies, S-11222) or streptavidin, Alexa Fluor 488 conjugate (Life Technologies, S-11223) at 1/250, and DAPI at 1/1000 in PBS were used.

### Primary DRG neuron imaging and analysis

Mounted cells were imaged using an Axioplan 2 microscope (Zeiss, Oberkochen, Germany), and single plane images analysed using ImageJ software (https://imagej.nih.gov/ij/). Cell bodies and processes of DRG neurons were visualised using the pan-neuronal marker *β*III-tubulin. To calculate the percentage of cells bearing neurites an average of 788 cells/coverslip were scored. Neuronal processes shorter than approximately half of the diameter of the cell body of the largest neurons (≈15 μm), as assessed by eye, were not counted as neurites. Neurite lengths were measured by manually plotting points along the length of a cell’s longest process, and an average of 51 cells/coverslip were assessed. Cell body area was measured by manually drawing around the circumference of the cell body, and an average of 202 cells/coverslip were assessed. To assess cleaved-caspase 3 staining, the average fluorescence intensity of *β*III-tubulin^+^ cell bodies was calculated for each coverslip, and the mean cell body fluorescence from the secondary only control was subtracted. DRG neurons were plated onto multiple coverslips, which were then PFA-fixed at different times post-plating (4, 48, and 96 h). Coverslips were simultaneously processed, and images acquired with the same confocal settings, so that fluorescence intensity could be compared across time points. An average of 57 cells/coverslip were assessed for cleaved-caspase 3 intensity. To categorise neurons, cells were divided into small (<315 μm^2^), medium (315-706 μm^2^), and large (>706 μm^2^) groups based on previously published criteria (30), and an average of 202 cells/coverslip were assessed. To assess the percentage of *β*III-tubulin^+^ cells expressing NF200, an average of 241 cells/coverslip were scored by eye. All DRG culture analyses include multiple coverslips (1-3)/animal and were performed blinded to genotype.

### Tissue imaging and analysis

All mounted tissues, except for spinal cord sections, were imaged using a LSM 780 laser scanning microscope (Zeiss), and images analysed using ImageJ software. To measure the area of *β*III-tubulin^+^ profiles in L1-L5 DRG sections, Z-stack images were taken with a 20x objective, and 3D projected (Max Intensity) images used to draw around the circumference of cell profiles. The areas of 300 profiles/DRG were averaged from three DRG to generate a single mean value for each mouse. Single plane, tile scan images of L1-L5 DRG sections were taken with a 10x or 20x objective in order to calculate the percentages of CGRP^+^, IB4^+^, NF200^+^, peripherin^+^, and parvalbumin^+^ cells. The Cell Counter ImageJ plugin was used to avoid recounting cells. An average of 470 profiles/DRG from three DRG were scored and used to calculate a mean value for each mouse. Cleaved-caspase 3 expression levels were assessed by measuring the average fluorescence intensity of each DRG section, and subtracting from this value the fluorescence of secondary-only control sections. All slides within a time point (i.e. P1, one, and three months) were processed in parallel and images acquired in the same session with identical settings. Three sections/DRG were used to produce a mean value for each DRG, which was then averaged across three DRG to get a cleaved-caspase 3 fluorescence intensity for each mouse. To perform *β*III-tubulin^+^ profile counts in L5 DRG, the volume of each DRG (in mm^3^) was estimated by calculating the mean area (in mm^2^) of 15-17 evenly-spaced sections and then multiplying this value by the section thickness and the number of sections taken for each DRG (71-116). The number of *β*III-tubulin^+^ profiles was counted in each of the 15-17 sections and averaged to get an estimate of neuron density (neurons/mm^3^). DRG volume and neuron density values were then used to estimate the number of profiles/L5 DRG. For soleus muscle spindle analyses, a full series of sections across the length of each muscle was used to count the number of spindles. Muscle spindles were identified based on SV2/2H3 fluorescence and, because this antibody combination is also used to visualise NMJs (64), stereotypical architecture. Spindle innervation status was assessed by eye by scoring the percentage of spindles displaying the characteristic circular SV2/2H3 fluorescence surrounding central, large circular nuclei (e.g. Fig. 3A, wild-type). Spindles lacking this staining around at least one nuclei were designated as not fully innervated (e.g. Fig. 3A, *Gars*^*C201R*/+^). All muscle spindles within each muscle were assessed. To analyse intraepidermal nerve fibre density, Z-stack images of the epidermis were taken of the central region of the lateral edge of glabrous hind paw using a 20x objective. Images were 3D projected (Max Intensity), uniformly thresholded across samples (PGP9.5 staining assigned to black and background to white), and all particles analysed within the epidermis. The summed pixel area was then divided by the epidermis area to get a percentage coverage of PGP9.5 staining/10 μm section. This value was halved for P1 samples, which were sectioned at twice the thickness of the one and three month plantar punches. Samples within time points were simultaneously processed and imaged with identical microscope settings. Six to eight sections were used to generate mean values for each animal, and secondary only control slides were used to ensure that background fluorescence did not contribute to the PGP9.5 coverage percentage. To assess E13.5 sensory nerve targeting of the hind limbs, single-plane, tile scan images were taken with a 20x objective. Images were then used to measure the distances between the ventral and dorsal main nerve trunk endings to the tips of the developing digits. The average distance of 2-6 nerves was calculated to produce values for each embryo. Nerve branching in the dorsal foot plate was also assessed by calculating the number of branches per mm of the longest length of each major nerve trunk and then averaging those values to produce a score for each embryo, similar to previously reported (65). In order to prevent subtle differences in developmental stage impacting the result, E13.5 statistical analyses were performed not on raw data but values relative to the wild-type mean.

Spinal cord sections were imaged with a LSM 700 laser scanning microscope (Zeiss), and analysed using ImageJ. Three tissue sections were assessed per animal, with four animals per genotype. All images were captured with a voxel size of 40 × 40 nm by 100 nm depth using a x63 Plan-Apochromat objective, suitable for deconvolution. Initially, a single image slice (≈75 μm width × 150 μm height) of IB4 and PSD95 labelling was taken across one randomly selected superficial dorsal horn, including laminae I-III, from the tissue section under analysis. Using the freehand drawing tool and ROI (Region of Interest) Manager in ImageJ and IB4 labelling as a marker of lamina II, this reference image was used to delineate regions of interest consisting of lamina I, II outer, II inner, and III. These regions of interest were used for assessing synapse densities across the different laminae of the dorsal horn. Sample image stacks were also captured of the same region, taken to a depth of 5.5 μm, and processed to derive binary representations of synaptic puncta. Firstly, stacks were deconvolved in ImageJ using the WPL deconvolution algorithm in the Parallel Iterative Deconvolution software package (66), based on the Iterative Deconvolve 3D plugin (67). 3D PSFs were generated in PSF Lab (68). Deconvolved image stacks were then filtered with the “despeckle” 3 × 3 median filter and thresholded using the OTSU method in ImageJ. To assess puncta within each region of interest, the image stack was divided into eight equally spaced image slices (0.1 μm z slices separated by 0.5 μm). Each region on the isolated slices was assessed separately for PSD95^+^ and synaptophysin^+^ puncta profiles using the Analyze Particles plugin in ImageJ. Puncta profile counts for each lamina were averaged across three tissue sections per animal, and plotted as puncta profile numbers per 100 μm^2^ of lamina. Phenotypic analyses of all tissues were performed blinded to genotype.

### Motor behaviour testing

Motor behaviour was assessed using a Grip Strength Meter (Bioseb, Vitrolles, France, GS3) and an accelerating Rota-Rod (Ugo Basile, Monvalle, Italy, 47600) as previously described (12, 69). Rota-Rod testing was sometimes completed before grip strength testing on the same day with at least a 30 min interval between tests. Mice were tested on two or three days within a week. Three trials were performed per animal per test day, with at least 5 min rest between trials. The maximum recorded value from the three trials in a day was used to calculate a mean value across days for each mouse.

### Sensory behaviour testing

For all sensory behaviour tests, mice were pre-acclimatised to the testing equipment and an average of 2-3 baseline readings were used to obtain a final measurement. To assess mechanosensation, Von Frey monofilaments were applied to the plantar surface of the hind paw and removal of the paw recorded as a positive response. A response threshold value was calculated using the up and down method (70). Proprioception was assessed using the beam walk test as previously described (71). Briefly, mice were filmed as they moved across a thin wooden beam of roughly one meter in length. The percentage of correct steps was calculated by recording the number of missed hind paw placements and hops, and comparing to the total number of steps taken for each run. Noxious mechanical thresholds were calculated with the use of an Analgesy-Meter (Ugo Basile, 37215) (72). Mice were lightly restrained and increasing mechanical pressure was applied either to the dorsum of the paw or to the base of the tail. The force at which an obvious withdrawal response was observed was taken as the withdrawal threshold. To assess heat sensitivity, the Hargreaves method was used (73). A radiant heat source was applied to the plantar surface of the hind paw and the latency to withdrawal was measured as the noxious heat threshold. All tests were carried out with the experimenter blind to genotype.

### Measurement and Analysis of Cytosolic Calcium

Cytosolic calcium in primary DRG neurons was assessed similar to previously described (74). Briefly, cells were washed twice with recording medium (156 mM NaCl, 10 mM D-glucose [Sigma, G8270], 10 mM HEPES [Sigma, H3375], 3 mM KCl [Sigma, P9333], 2 mM CaCl_2_ [Sigma, C8106], 2 mM MgSO_4_ [Sigma, M7506], and 1.25 mM KH_2_PO_4_, pH 7.34-7.36 - adjusted with 10 M NaOH [Sigma, 72068]), and loaded with 5 μM fura-2 AM (Life Technologies, F1221) and 0.002% (v/v) pluronic acid F-127 (Life Technologies, P3000MP) in recording medium for 30 min at room temperature. Fura-2 was subsequently removed and cells washed twice in recording medium before cytosolic calcium was detected in single cells every 2 s using an IX70 inverted microscope (Olympus, Tokyo, Japan), a cooled Retiga Fast 1394 CCD camera (QImaging, Surrey, Canada), and Andor iQ 1.9 software (Belfast, UK). Following at least 90 s continuous recording of the baseline fura-2 ratio, the medium was carefully replaced with recording medium containing either 50 mM KCl or 1 μM capsaicin (Sigma, M2028). To calculate the fura-2 ratio change from baseline to maximal response for individual cells, the mean of 20 sequential recordings post-stimulation was divided by the mean of 20 sequential baseline recordings prior to stimulation. The mean average was then calculated for all cells simultaneously imaged in a well. An average of 10 cells per well was assessed. Capsaicin responsive and non-responsive cells were categorised based on whether the percentage fura-2 ratio change was ≥ 150% or < 150%, respectively. Fura-2 experiments include multiple wells (2-3)/treatment/animal.

### Statistical analysis

Data were assumed to be normally distributed unless evidence to the contrary could be provided by the D’Agostino and Pearson omnibus normality test. Data were statistically analysed using an unpaired *t*-test, or one- or a two-way analysis of variance (ANOVA) with Sidak’s multiple comparisons tests. If the data did not pass normality testing, Mann-Whitney *U* tests or Kruskal-Wallis tests with Dunn’s multiple comparison tests were used. GraphPad Prism 6 software was used for all statistical analyses and production of figures. Means ± standard error of the mean (S.E.M.) are plotted for all graphs.

## Acknowledgements

The authors would like to thank members of the Schiavo, Greensmith (Institute of Neurology, UCL), Talbot, Cader, and Bennett (University of Oxford) laboratories for productive discussions, Andrey Y. Abramov, J. Barney Bryson, Benjamin E. Clarke, Steven Middleton, Gustavo Pregoni, Annina B. Schmid, Greg A. Weir, and Emma R. Wilson for sharing experimental expertise, Nathalie Schmieg for providing the E18.5 *Kidins220*^-/-^ brain, and Alexander M. Rossor for critical reading of the manuscript.

Funding

This work was supported by a Wellcome Trust Sir Henry Wellcome Postdoctoral Fellowship (103191/Z/13/Z) [JNS], the French Muscular Dystrophy Association (AFM-Telethon) [JNS, MZC, KT], a Wellcome Trust Senior Investigator Award (107116/Z/15/Z) [GS], University College London [GS], a UK Medical Research Council research grant [JMD], and the National Institutes of Health (NS054154) [RWB].

## Author Contributions

JNS conceived the work; JNS, JMD, SJW, DLB, and GS designed the experiments; JNS, JMD, SJW, ELS, and AG-M performed the experiments, JNS, JMD, and SJW analysed the data; all authors contributed to the writing of the paper and have approved submission of this work. The funders had no role in study design, data collection and analysis, decision to publish, or preparation of the manuscript.

## Supplementary Figures

**Supplementary Figure 1.**
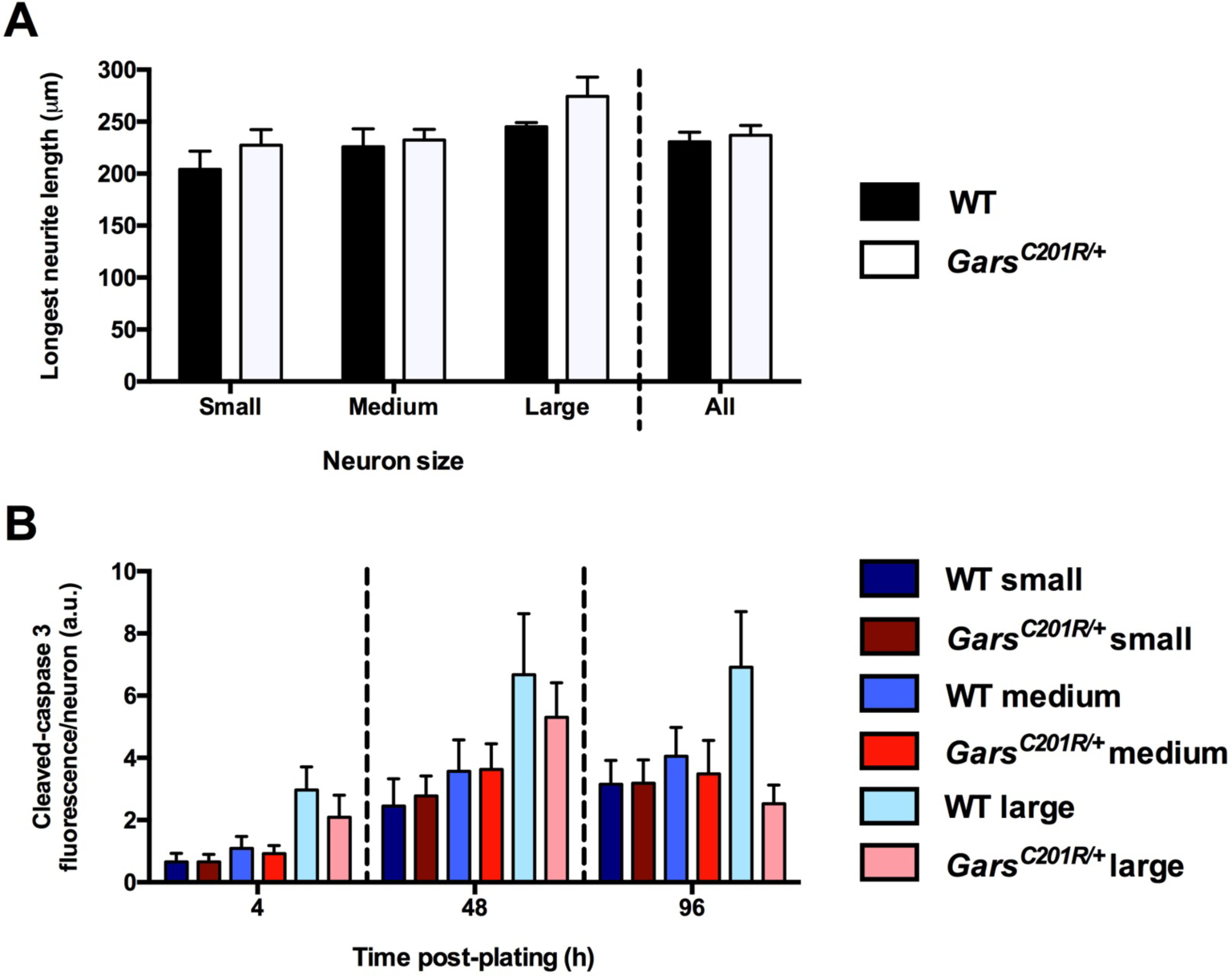
Different classes of mutant sensory neuron, based on cell soma size, are equally unaffected. **(A)** Small (<315 μm^2^), medium (315-706 μm^2^), and large (>706 μm^2^) area sensory neurons show no difference in the longest neurite length between wild-type and *Gars*^*C201R*/+^ cultures. Neurons were stained with anti-*β*III-tubulin. *P* = 0.05, Kruskal-Wallis test, *P* > 0.05, Dunn’s multiple comparison tests between wild-type and mutant samples of all cell sizes (excluding “All” category, which was tested in Fig. 1B, top right). **(B)** There is also no evidence for greater cell death, as assessed by cleaved-caspase 3 fluorescence intensity per neuron. 4 h time point, *P* = 0.047, Kruskal-Wallis test, P > 0.05, Dunn’s multiple comparison tests between wild-type and mutant samples of all cell sizes; 48 h time point, two-way ANOVA (*P* = 0.018, cell body size; *P* = 0.729, genotype; *P* = 0.739, interaction); 96 h time point, two-way ANOVA (*P* = 0.362, cell body size; *P* = 0.071, genotype; *P* = 0.099, interaction). a.u., arbitrary units. Four mice/genotype were analysed.

**Supplementary Figure 2.**
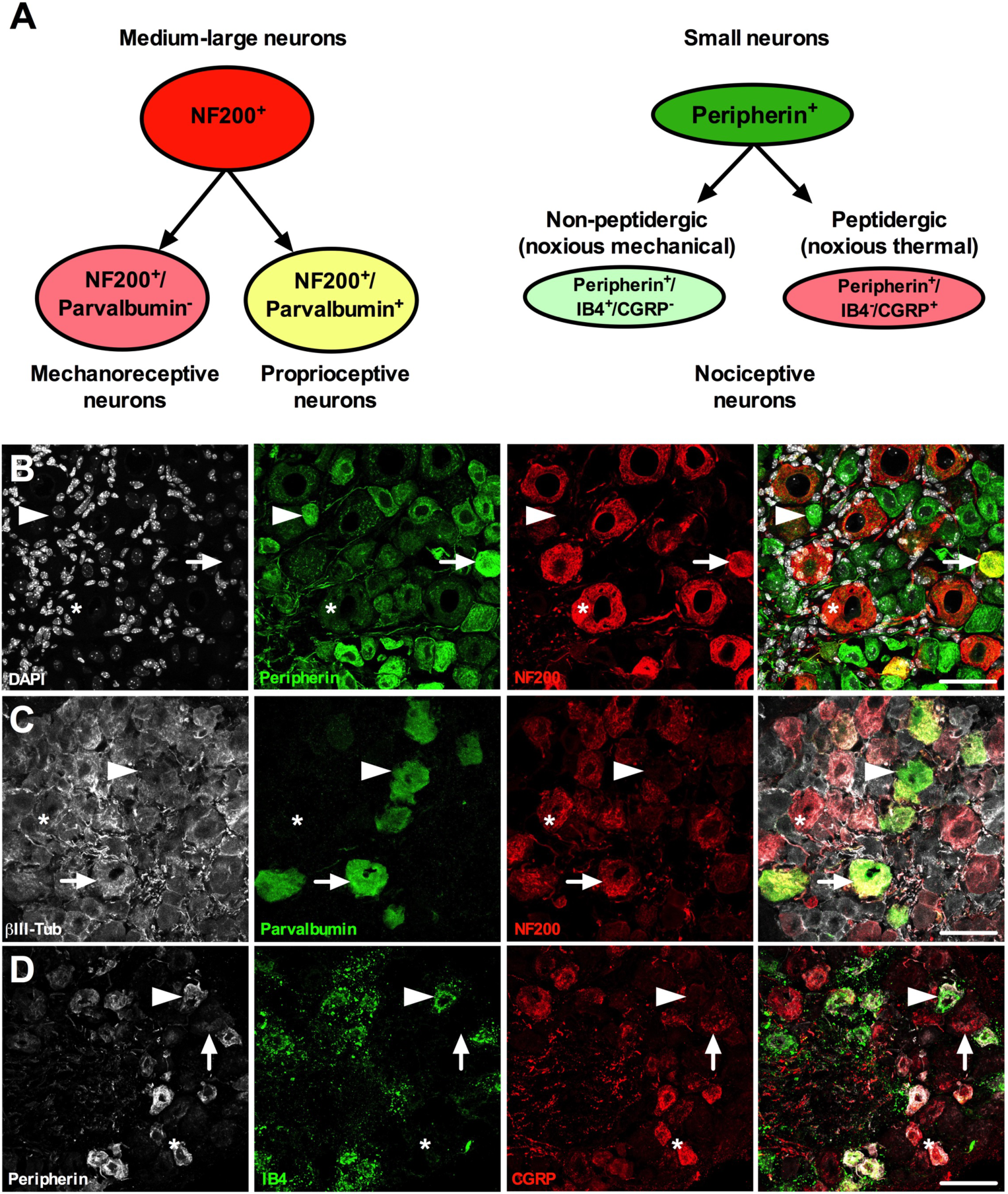
Immunohistochemical analysis of functional sensory neuron subtype markers. **(A)** Different functional classes of sensory neurons can be identified based on the expression of marker proteins. NF200 and peripherin are mostly exclusively expressed in medium-large (A-fibres) and small (Aδ and C-fibres) sensory nerves, respectively. Those NF200^+^ cells that lack parvalbumin expression are considered to be mechanoreceptive neurons, while those that are parvalbumin^+^ are proprioceptive (A, left-hand side). Peripherin^+^ cells that co-stain with isolectin B4 (IB4), but not calcitonin gene-related peptide (CGRP), are non-peptidergic, mainly mechanical nociceptors, and those peripherin+ cells that are labelled with CGRP, but not IB4, are peptidergic, principally thermal nociceptors (A, right-hand side). **(B)** Representative example of co-staining with DAPI (white), peripherin (green), and NF200 (red). The asterisk highlights a NF200^+^/peripherin^−^ medium-large neuron, and the arrowhead marks a NF200^−^/peripherin^+^ small neuron. The arrow identifies an infrequently seen (≈ 2-3%) cell type that expresses both NF200 and peripherin. A second such cell (yellow) can be seen towards the bottom of the merged image. **(C)** Representative example of co-staining with *β*III-tubulin (white), parvalbumin (green), and NF200 (red). The asterisk identifies a NF200^+^/parvalbumin^−^ cell likely to be a mechanoreceptive sensory neuron, the arrow marks a NF200^+^/parvalbumin^+^ proprioceptive nerve, and the arrowhead highlights a rare NF200^−^/parvalbumin^+^ cell. **(D)** Representative example of co-staining with peripherin (white), IB4 (green), and CGRP (red). The arrowhead identifies a peripherin^+^/IB4^+^/CGRP^−^ neuron, which is likely a mechanosensitive nociceptor, the asterisk marks a peripherin^+^/IB4^−^/CGRP^+^ nociceptor probably sensitive to noxious thermal stimuli, and the arrow highlights an uncommon (≈ 1%) example of a peripherin^−^/IB4^−^/CGRP^+^ cell. All images are collapsed Z-stacks taken of one month wild-type DRG. Scale bars = 50 μm.

**Supplementary Figure 3.**
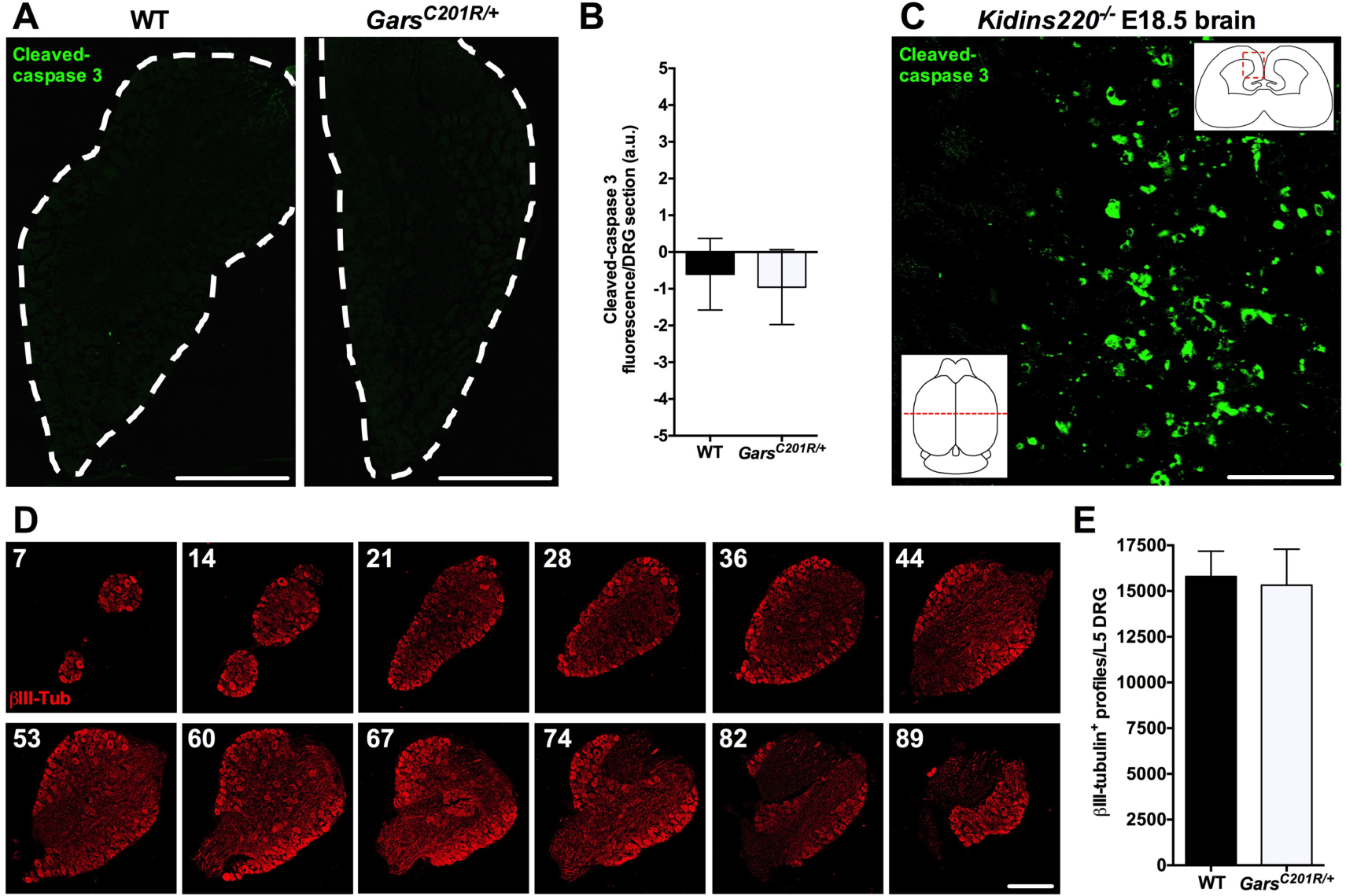
*Gars*^*C201R*/+^ DRG show no signs of sensory neuron loss at one month. **(A)** Representative single confocal plane images of one month old wild-type (left) and *Gars*^*C201R*/+^ (right) DRG sections stained for the apoptotic marker cleaved-caspase 3. DRG sections are outlined by dashed lines. **(B)** There is no difference in cleaved-caspase 3 fluorescence intensity between wild-type and mutant DRG. *P* = 0.812, unpaired *t*-test. a.u., arbitrary units. **(C)** Representative cleaved-caspase 3 staining of a section taken from an E18.5 *Kidins220*^−/-^ brain, which was used as a positive control. Inset images depict the region of the coronal section (bottom left), and the area of the section that was imaged (top right). **(D)** Representative images of serial sections taken from a single wild-type L5 DRG and stained with *β*III-tubulin (red). The number in the top left of each panel depicts the section number. Serial sections were taken throughout the entire length of the DRG in order to estimate the number of sensory neurons per L5 ganglion. **(E)** *β*III-tubulin cell profile estimates indicate that *Gars*^*C201R*/+^ L5 DRG show no signs of sensory neuron loss. *P* = 0.849, unpaired *t*-test. Four mice/genotype were analysed. All images are single confocal planes. Scale bars = 200 μm (A, D) and 50 μm (C).

**Supplementary Figure 4.**
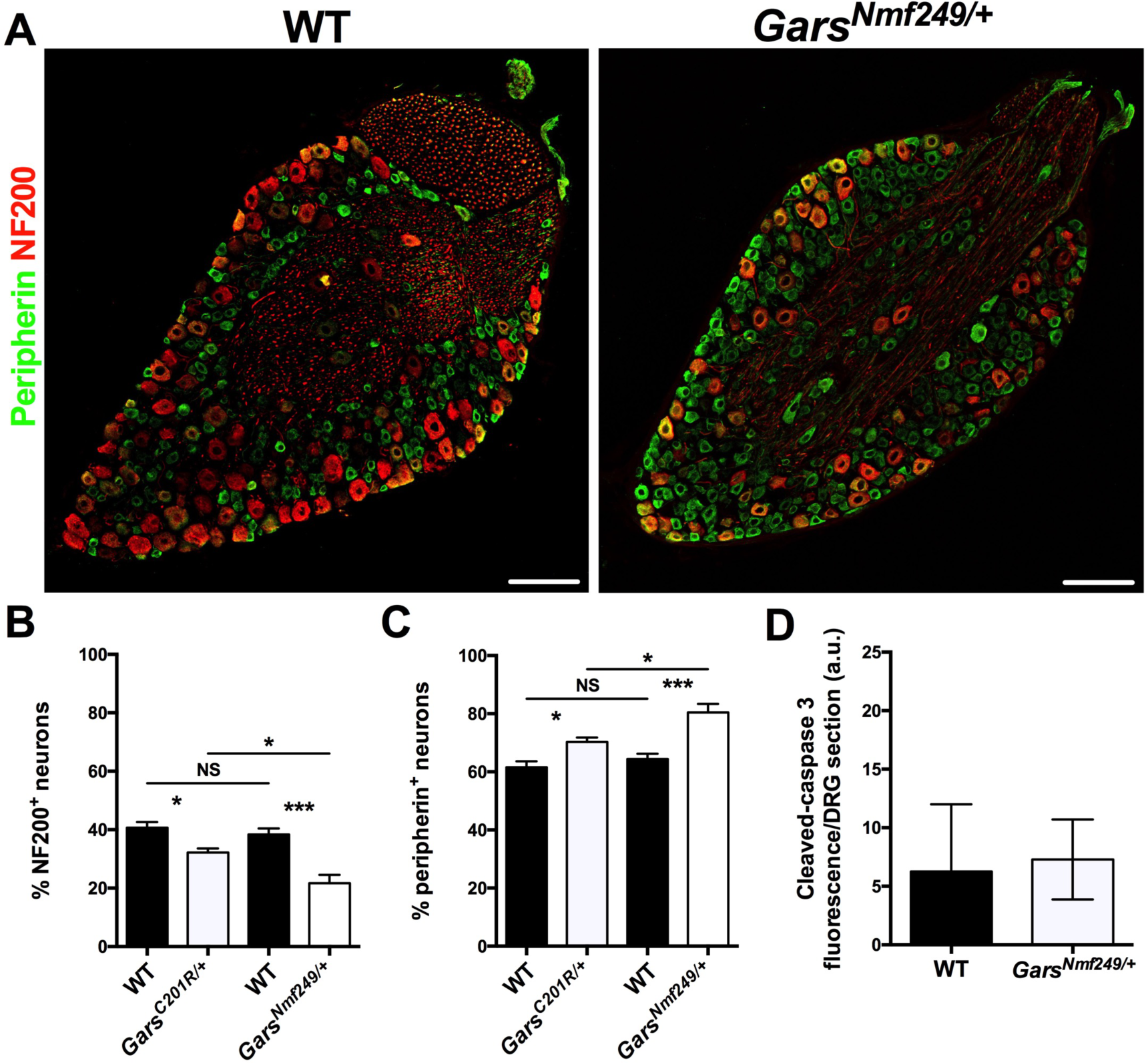
*Gars* mouse DRG defects correlate with mutant severity. **(A)** Representative single confocal plane images of one month old DRG taken from wild-type (left) and the more severe CMT2D mutant mouse model, *Gars*^*Nmf249*/+^, stained for peripherin (green) and NF200 (red). Scale bars = 100 μm. **(B-C)** *Gars*^*Nmf249*/+^ mice display a significant reduction in the percentage of NF200^+^ cells (B, *P* < 0.001, one-way ANOVA), and a concomitant decrease in the percentage of peripherin^+^ cells (C, *P* < 0.001, one-way ANOVA) compared to both wild-type and *Gars*^*C201R*/+^ mice. NS, not significant, * *P* < 0.05, *** *P* < 0.001, Sidak’s multiple comparisons test. *Gars*^*C201R*/+^ data are taken from Fig. 2. **(D)** There is no evidence for increased levels of cell death in the *Gars*^*Nmf249*/+^ DRG, as assessed by activated caspase 3 staining. a.u., arbitrary units. *P* = 0.882, unpaired *t*-test.

**Supplementary Figure 5.**
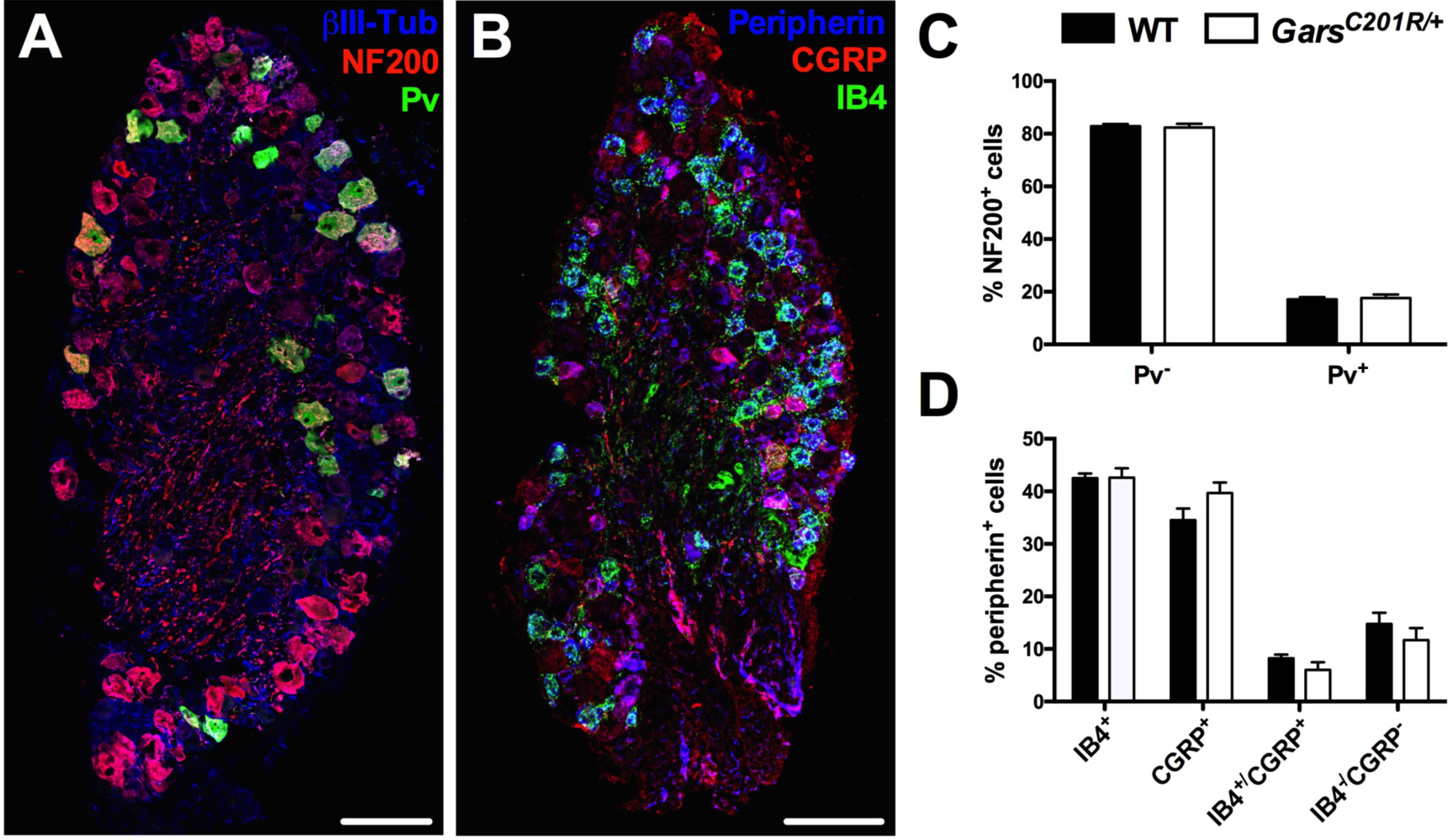
*Gars*^*C201R*/+^ mechanoreceptive and proprioceptive sensory neurons are equally affected, as are nociceptor subtypes. **(A-B)** Representative DRG sections from a one month old wild-type mouse stained to identify mechanoreceptive neurons (A, NF200^+^ [red]/Pv^−^), proprioceptive neurons (A, NF200^+^/Pv^+^[green]), non-peptidergic nociceptors (B, peripherin^+^[blue]/IB4^+^[green]/CGRP^−^), and peptidergic nociceptors (B, peripherin^+^/IB4^−^/CGRP^+^[red]). Images are single confocal planes. Scale bars = 100 μm. **(C)** *Gars*^*C201R*/+^ DRG show no difference in the percentage of NF200^+^ cells that co-stain for the proprioceptive marker parvalbumin (Pv). *P* = 0.768, unpaired *t*-test between Pv^−^ cells. **(D)** There is also no difference between the percentages of wild-type and mutant peripherin^+^ sensory neurons expressing either IB4 or CGRP. *P* = 0.964 and *P* = 0.132, unpaired *t*-test between IB4^+^ cells and CGRP^+^ cells, respectively. Four mice/genotype were analysed.

**Supplementary Figure 6.**
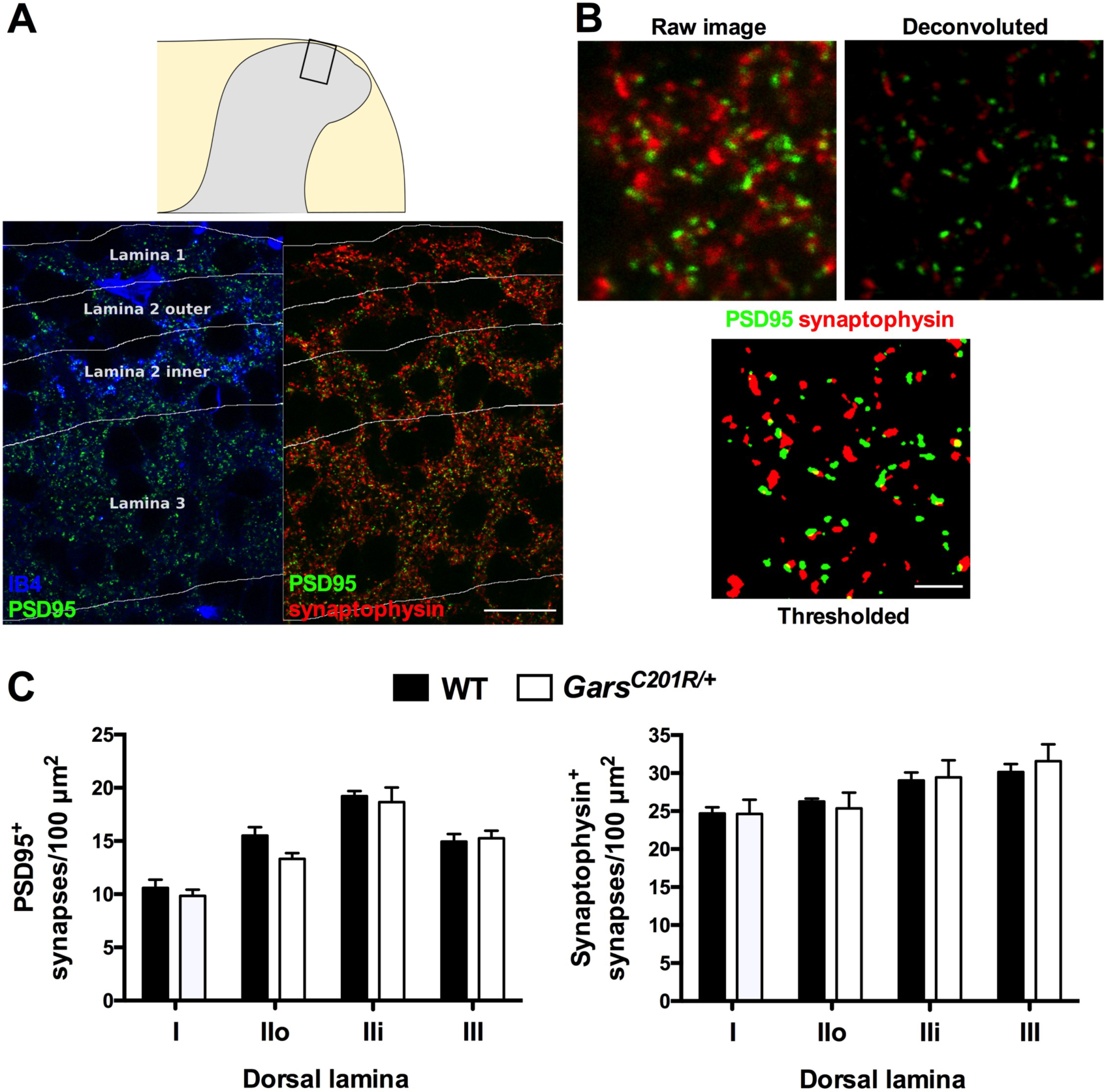
Synaptic densities in dorsal horn laminae I-III remain unchanged in one month old *Gars* mice. **(A)** Representative collapsed Z-stack image of a spinal cord section from a one month old wild-type mouse stained for the post-synaptic marker PSD95 (green, bottom), the pre-synaptic protein synaptophysin (red, bottom right), and IB4 (blue, bottom left) to identify lamina II and delineate regions of interest. The schematic (top) depicts the region of the dorsal horn that was imaged. **(B)** Representative images of the deconvolution and thresholding processes used to analyse synapse numbers in distinct laminae (see methods). **(C)** There is no difference in synapse density in dorsal horn laminae I-III between wild-type and *Gars*^*C201R*/+^ mice when using either PSD95 (two-way ANOVA, *P* < 0.001, lamina; *P* = 0.170, genotype; *P* = 0.472, interaction) or synaptophysin (two-way ANOVA, *P* = 0.002, lamina; *P* = 0.847, genotype; *P* = 0.908, interaction) to assess synapse numbers. Intra-laminae comparisons between genotypes also show no difference (*P* > 0.05, Sidak’s multiple comparisons test). IIo and IIi, outer and inner regions of lamina II, respectively. Four mice/genotype were analysed. Scale bars = 20 μm (A) and 2 μm (B).

**Supplementary Figure 7.**
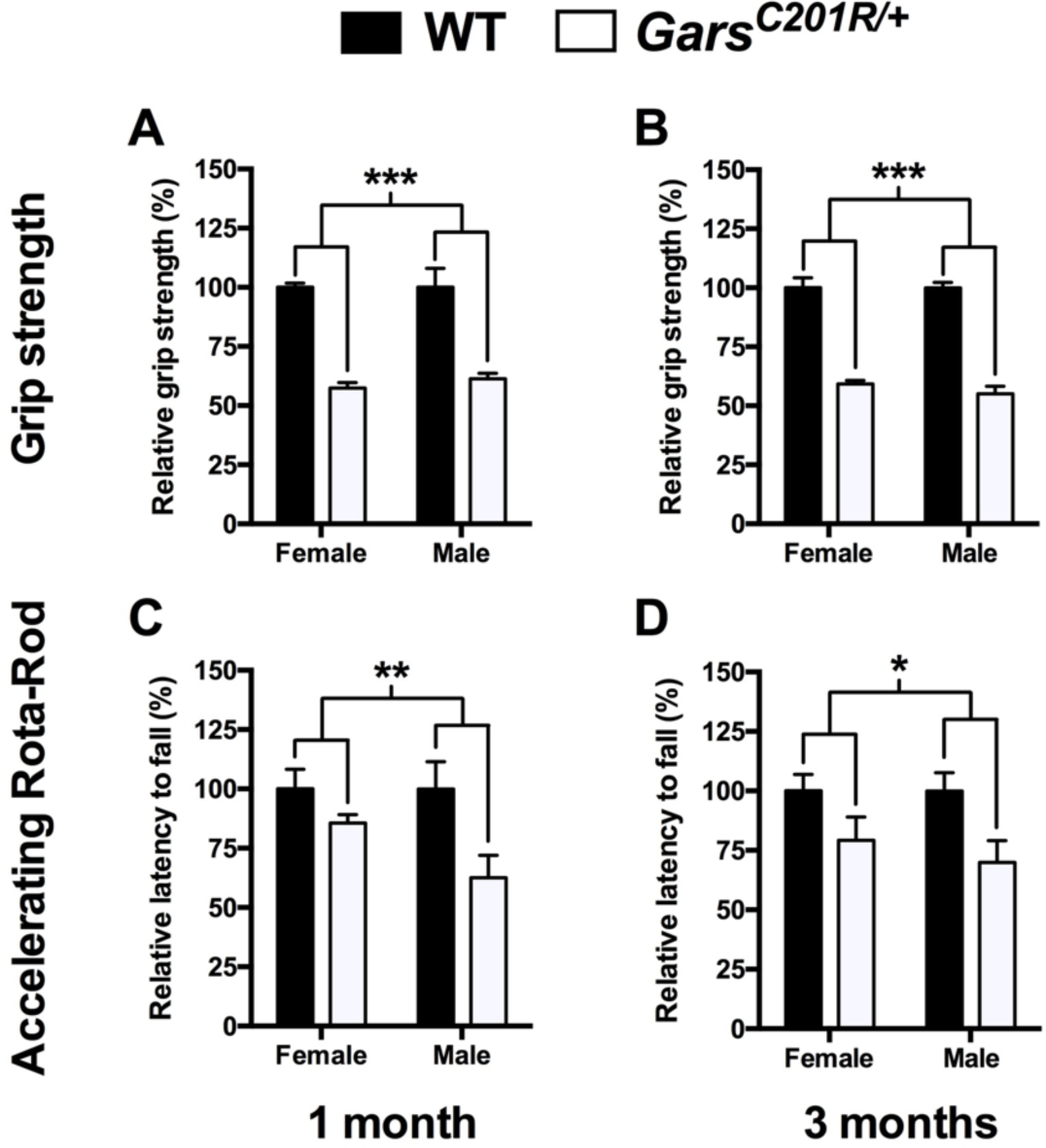
*Gars*^*C201R*/+^ mice display motor defects that remain relatively stable from one to three months. **(A-B)** Both female and male *Gars*^*C201R*/+^ mice show a significant reduction in grip strength compared to wild-type mice of the same sex at one (A, two-way ANOVA, *P* = 0.854, sex; *** *P* < 0.001, genotype; *P* = 0.616, interaction) and three (B, two-way ANOVA, *P* < 0.001, sex; *** *P* < 0.001, genotype; *P* = 0.037, interaction) months. **(C-D)** Similarly, *Gars*^*C201R*/+^ mice show a significant reduction in the time taken to fall off an accelerating Rota-Rod at one (C, two-way ANOVA, *P* = 0.326, sex; ** *P* = 0.007, genotype; *P* = 0.188, interaction) and three months (D, two-way ANOVA, *P* < 0.001, sex; * *P* = 0.010, genotype; *P* = 0.972, interaction). In both behavioural tests, the mutant defects remained relatively stable from one to three months (female grip strength, *P* = 0.523; male grip strength, *P* = 0.155; female Rota-Rod, *P* = 0.556; male Rota-Rod, *P* = 0.590, unpaired *t*-test). Six mice/genotype/sex were analysed. The statistical tests of significance represented on the figures were performed on raw data (Supplementary Tables 8 and 9), while the percentages relative to wild-type, which are plotted, were used to compare mutant progression over time.

**Supplementary Figure 8.**
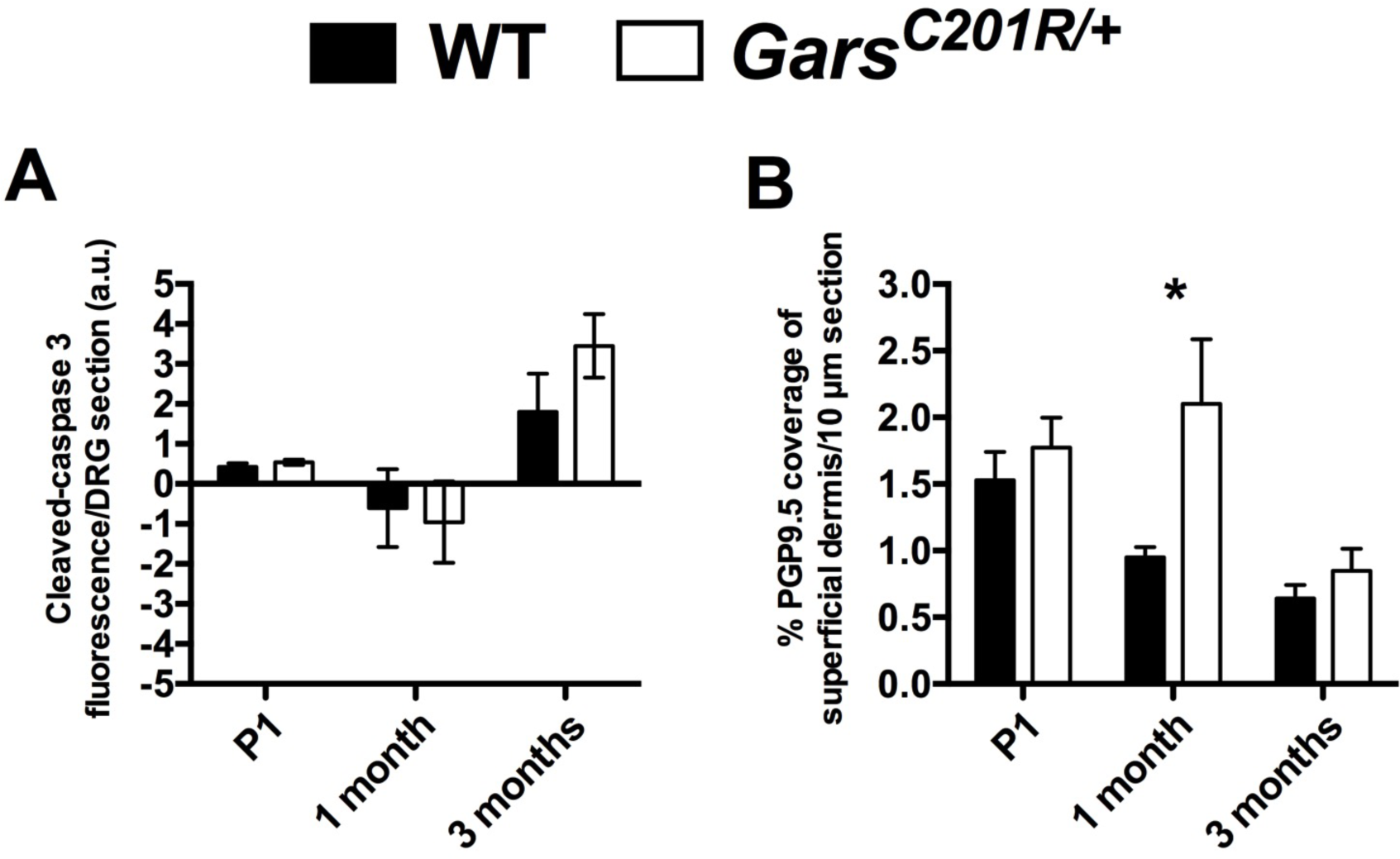
Longitudinal analysis of cell death in lumbar DRG and intraepidermal nerve fibre density in glabrous hind paw. **(A)** There is no difference in the cleaved-caspase 3 fluorescence intensity per neuron between wild-type and *Gars*^*C201R*/+^ DRG at any time point tested, suggesting that cell death is not accounting for the observed DRG cellular phenotype. Two-way ANOVA (*P* < 0.001, age; *P* = 0.444, genotype; *P* = 0.413, interaction). **(B)** PGP9.5 staining in the hind paw epidermis of wild-type mice decreases progressively from P1 to three months. *Gars*^*C201R*/+^ mice show no difference at P1, significantly more innervation at one month, but then no difference again at three months. Two-way ANOVA (*P* = 0.002, age; *P* = 0.013, genotype; *P* = 0.132, interaction). * *P* < 0.05, ** *P* < 0.01, Sidak’s multiple comparisons test. 3-5 (A) and 4-6 (B) mice/genotype/time point were analysed. Data from the one month time point in panels A and B are taken from Fig. S3B and Fig. 3E, respectively.

## Supplementary Tables

**Supplementary Table 1.** Primary antibodies successfully used in the study. C-C3, cleaved-caspase 3; CST, Cell Signalling Technology; DSHB, Developmental Studies Hybridoma Bank; ELS, Enzo Life Sciences; FI, Frontier Institute; M, monoclonal; P, polyclonal; Syp, synaptophysin; SV2, synaptic vesicle 2; 2H3, neurofilament. For failed staining, the highest concentration is reported.

| Antigen | Source<br>(Product #) | Host species<br>(clonality) | Concentration used |  |  |  |  |  |  |
| --- | --- | --- | --- | --- | --- | --- | --- | --- | --- |
|  |  |  | DRG<br>cultures | DRG | Skin<br>axons | Muscle<br>spindles | Spinal<br>cord | E18.5<br>brain | E13.5<br>feet |
| CGRP | ELS<br>(BML-CA1137) | Sheep (P) | - | 1/200 | - | - | - | - | - |
| C-C3 | CST (9661) | Rabbit (P) | 1/500 | 1/500 | - | - | - | 1/500 | - |
| Laminin | Sigma<br>(L9393) | Rabbit (P) | - | - | - | 1/1000 | - | - | - |
| NF200 | Sigma<br>(N0142) | Mouse (M) | 1/500 | 1/500 | Failed at<br>1/500 | - | - | - | - |
| Parvalbumin | Swant<br>(PV 27) | Rabbit<br>(unknown) | - | 1/1000 | - | Failed at<br>1/500 | - | - | - |
| Peripherin | Merck<br>(AB1530) | Rabbit (P) | Failed at<br>1/500 | 1/500 | Failed at<br>1/500 | - | - | - | - |
| PGP9.5 | UltraClone<br>(RA 95101) | Rabbit (P) | - | - | 1/1000 | - | - | - | - |
| PSD95 | FI<br>(Af628) | Rabbit (P) | - | - | - | - | 1/200 | - | - |
| SV2 | DSHB<br>(concentrate) | Mouse (M) | - | - | - | 1/250 | - | - | - |
| Syp | FI<br>(Af300) | Guinea Pig<br>(P) | - | - | - | - | 1/200 | - | - |
| βIII-tubulin | Abcam<br>(ab41489) | Chicken (P) | 1/500 | 1/500 | - | - | - | - | - |
| βIII-tubulin | Covance<br>(mms-435p) | Mouse (M) | - | 1/500 | - | - | - | - | - |
| 2H3 | DSHB<br>(supernatant) | Mouse (M) | - | - | - | 1/250 | - | - | 1/50 |

**Supplementary Table 2.**
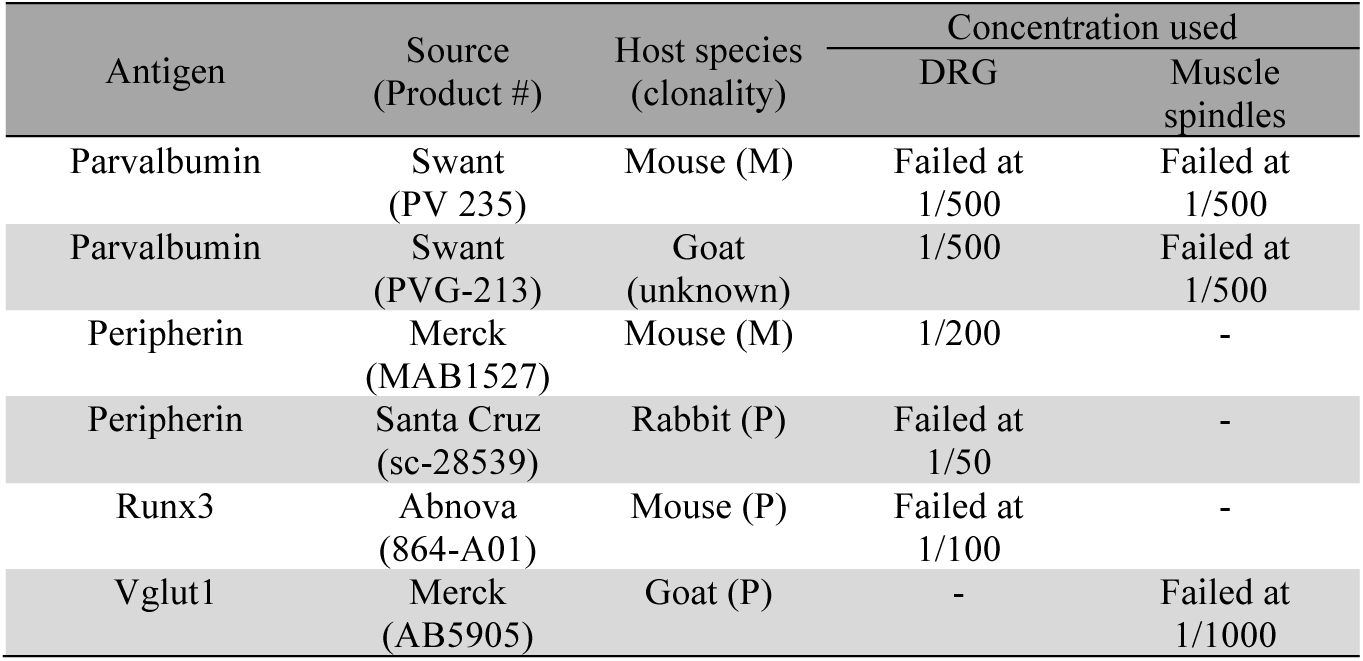
Additional primary antibodies tested in the study. M, monoclonal; P, polyclonal. For failed staining, the highest concentration tested is reported.

| Antigen | Source<br>(Product #) | Host species<br>(clonality) | Concentration used |  |
| --- | --- | --- | --- | --- |
|  |  |  | DRG | Muscle<br>spindles |
| Parvalbumin | Swant<br>(PV 235) | Mouse (M) | Failed at<br>1/500 | Failed at<br>1/500 |
| Parvalbumin | Swant<br>(PVG-213) | Goat<br>(unknown) | 1/500 | Failed at<br>1/500 |
| Peripherin | Merck<br>(MAB1527) | Mouse (M) | 1/200 | - |
| Peripherin | Santa Cruz<br>(sc-28539) | Rabbit (P) | Failed at<br>1/50 | - |
| Runx3 | Abnova<br>(864-A01) | Mouse (P) | Failed at<br>1/100 | - |
| Vglut1 | Merck<br>(AB5905) | Goat (P) | - | Failed at<br>1/1000 |

**Supplementary Table 3.** Raw data generated by the Von Frey test of mechanosensation at one and three months. *** *P* < 0.001, compared to wild-type using Sidak’s multiple comparison test.

| Age (months) | Von Frey withdrawal threshold (g) |  |
| --- | --- | --- |
|  | Wild-type | *Gars*<sup>C201R/+</sup> |
| 1 | 0.38 ± 0.028 | 0.48 ± 0.035 |
| 3 | 0.49 ± 0.035 | 0.66 ± 0.041** |

**Supplementary Table 4.**
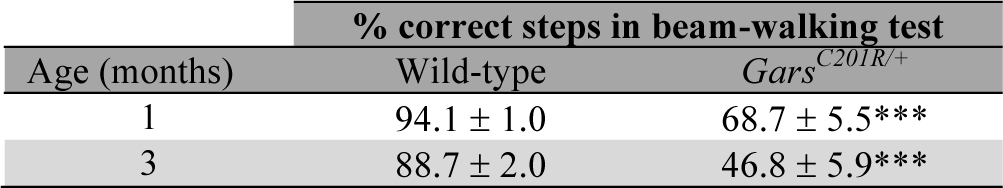
Raw data generated by the beam-walking test of proprioception at one and three months. *** *P* < 0.001, compared to wild-type using Dunn’s multiple comparison test.

**Supplementary Table 5.** Raw data generated by the Randall-Selitto test of hind paw mechanical nociception at one and three months. *** *P* < 0.001, compared to wild-type using Sidak’s multiple comparisons test.

| Age (months) | Randall-Selitto paw threshold (g) |  |
| --- | --- | --- |
|  | Wild-type | *Gars*<sup>C201R/+</sup> |
| 1 | 123.3 ± 9.6 | 76.5 ± 5.9*** |
| 3 | 135.2 ± 9.9 | 74.3 ± 3.6*** |

**Supplementary Table 6.** Raw data generated by the Randall-Selitto test of tail mechanical nociception at one and three months. ** *P* < 0.01, *** *P* < 0.001, compared to wild-type using Sidak’s multiple comparisons test.

| Age (months) | Randall-Selitto tail threshold (g) |  |
| --- | --- | --- |
|  | Wild-type | *Gars*<sup>C201R/+</sup> |
| 1 | 104.1 ± 15.3 | 59.6 ± 4.6** |
| 3 | 153.0 ± 9.6 | 81.9 ± 5.2*** |

**Supplementary Table 7.** Raw data generated by the Hargreaves test of thermal nociception at one and three months. * *P* < 0.05, *** *P* < 0.001, compared to wild-type using Sidak’s multiple comparisons test.

| Age (months) | Hargreaves latency to withdrawal (s) |  |
| --- | --- | --- |
|  | Wild-type | *Gars*<sup>C201R/+</sup> |
| 1 | 7.06 ± 0.53 | 5.44 ± 0.46* |
| 3 | 9.03 ± 0.53 | 5.84 ± 0.41*** |

**Supplementary Table 8.**
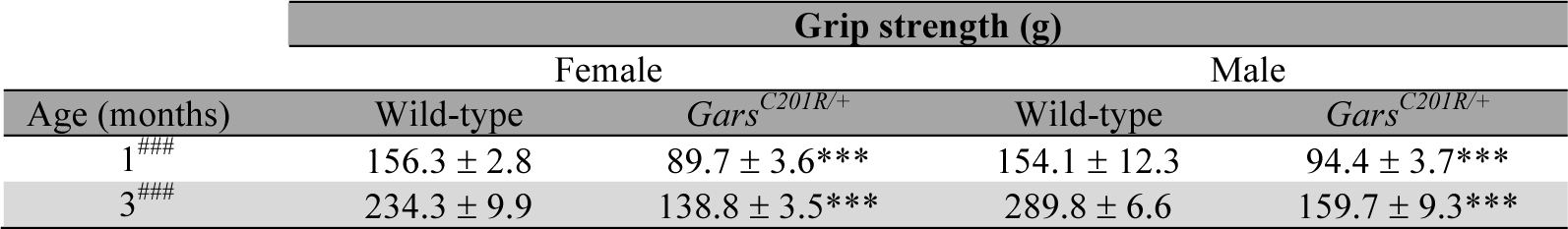
Raw data generated by the grip strength test at one and three months. *** *P* < 0.001, compared to wild-type using Sidak’s multiple comparisons test. ### *P* < 0.001, two-way ANOVA comparison of genotypes.

**Supplementary Table 9.**
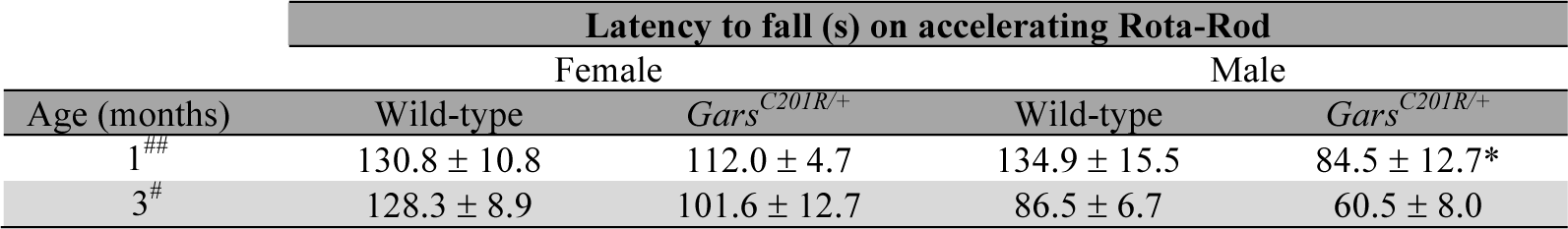
Raw data generated by the accelerating Rota-Rod test at one and three months. * *P* < 0.05, compared to wild-type using Sidak’s multiple comparisons test. # *P* < 0.05, ## *P* < 0.01, two-way ANOVA comparison of genotypes.

## References

1. Reilly MM, Murphy SM, Laura M (2011) Charcot-Marie-Tooth disease. J Peripher Nerv Syst 16(1): 1–14.

2. El-Abassi R, England JD, Carter GT (2014) Charcot-Marie-Tooth disease: an overview of genotypes, phenotypes, and clinical management strategies. PM R 6(4):342–355.

3. Skre H (1974) Genetic and clinical aspects of Charcot-Marie-Tooth’s disease. Clin Genet 6(2):98–118.

4. Antonellis A, et al. (2003) Glycyl tRNA synthetase mutations in Charcot-Marie-Tooth disease type 2D and distal spinal muscular atrophy type V. Am J Hum Genet 72(5):1293–1299.

5. Jordanova A, et al. (2006) Disrupted function and axonal distribution of mutant tyrosyl-tRNA synthetase in dominant intermediate Charcot-Marie-Tooth neuropathy. Nat Genet 38(2):197–202.

6. Latour P, et al. (2010) A major determinant for binding and aminoacylation of tRNA^Ala^ in cytoplasmic Alanyl-tRNA synthetase is mutated in dominant axonal Charcot-Marie-Tooth disease. Am J Hum Genet 86(1):77–82.

7. McLaughlin HM, et al. (2010) Compound heterozygosity for loss-of-function lysyl-tRNA synthetase mutations in a patient with peripheral neuropathy. Am J Hum Genet 87(4):560–566.

8. Vester A, et al. (2013) A loss-of-function variant in the human histidyl-tRNA synthetase (*HARS*) gene is neurotoxic *in vivo*. Hum Mut 34(1): 191–199.

9. Motley WW, Talbot K, Fischbeck KH (2010) *GARS* axonopathy: not every neuron’s cup of tRNA. Trends Neurosci 33(2):59–66.

10. Storkebaum E (2016) Peripheral neuropathy via mutant tRNA synthetases: Inhibition of protein translation provides a possible explanation. Bioessays (in press) 10.1002/bies.201600052.

11. Seburn KL, Nangle LA, Cox GA, Schimmel P, Burgess RW (2006) An active dominant mutation of glycyl-tRNA synthetase causes neuropathy in a Charcot-Marie-Tooth 2D mouse model. Neuron 51(6):715–726.

12. Achilli F, et al. (2009) An ENU-induced mutation in mouse glycyl-tRNA synthetase (*GARS*) causes peripheral sensory and motor phenotypes creating a model of Charcot-Marie-Tooth type 2D peripheral neuropathy. Dis Model Mech 2(7-8):359–373.

13. Nangle LA, Zhang W, Xie W, Yang XL, Schimmel P (2007) Charcot-Marie-Tooth disease-associated mutant tRNA synthetases linked to altered dimer interface and neurite distribution defect. Proc Natl Acad Sci U S A 104(27): 11239–11244.

14. Xie W, Nangle LA, Zhang W, Schimmel P, Yang XL (2007) Long-range structural effects of a Charcot-Marie-Tooth disease-causing mutation in human glycyl-tRNA synthetase. Proc Natl Acad Sci U S A 104(24): 9976–9981.

15. Motley WW, et al. (2011) Charcot-Marie-Tooth-linked mutant GARS is toxic to peripheral neurons independent of wild-type GARS levels. PLoS Genet 7(12):e1002399.

16. He W, et al. (2011) Dispersed disease-causing neomorphic mutations on a single protein promote the same localized conformational opening. Proc Natl Acad Sci USA 108(30):12307–12312.

17. Grice SJ, et al. (2015) Dominant, toxic gain-of-function mutations in gars lead to non-cell autonomous neuropathology. Hum Mol Genet 24(15):4397–4406.

18. He W, et al. (2015) CMT2D neuropathy is linked to the neomorphic binding activity of glycyl-tRNA synthetase. Nature 526(7575):710–714.

19. Sivakumar K, et al. (2005) Phenotypic spectrum of disorders associated with glycyl-tRNA synthetase mutations. Brain 128(Pt 10):2304–2314.

20. Del Bo R, et al. (2006) Coexistence of CMT-2D and distal SMA-V phenotypes in an Italian family with a GARS gene mutation. Neurology 66(5):752–754.

21. Hamaguchi A, Ishida C, Iwasa K, Abe A, Yamada M (2010) Charcot-Marie-Tooth disease type 2D with a novel glycyl-tRNA synthetase gene (GARS) mutation. J Neurol 257(7): 1202–1204.

22. Sun A, et al. (2015) A novel mutation of the glycyl-tRNA synthetase (GARS) gene associated with Charcot-Marie-Tooth type 2D in a Chinese family. Neurol Res 37(9):782–787.

23. Lee HJ, et al. (2012) Two novel mutations of GARS in Korean families with distal hereditary motor neuropathy type V. J Peripher Nerv Syst 17(4):418–421.

24. Le Pichon CE, Chesler AT (2014) The functional and anatomical dissection of somatosensory subpopulations using mouse genetics. Front Neuroanat 8:21.

25. Lallemend F, Ernfors P (2012) Molecular interactions underlying the specification of sensory neurons. Trends Neurosci 35(6):373–381.

26. Sleigh JN, Grice SJ, Burgess RW, Talbot K, Cader MZ (2014) Neuromuscular junction maturation defects precede impaired lower motor neuron connectivity in Charcot-Marie-Tooth type 2D mice. Hum Mol Genet 23(10):2639–2650.

27. Harper AA, Lawson SN (1985) Conduction velocity is related to morphological cell type in rat dorsal root ganglion neurones. J Physiol 359:31–46.

28. Chen XJ, et al. (2007) Proprioceptive sensory neuropathy in mice with a mutation in the cytoplasmic Dynein heavy chain 1 gene. J Neurosci 27(52):14515–14524.

29. Sassone J, et al. (2016) ALS mouse model SOD1^G93A^ displays early pathology of sensory small fibers associated to accumulation of a neurotoxic splice variant of peripherin. Hum Mol Genet 25(8):1588–99.

30. Sommer EW, Kazimierczak J, Droz B (1985) Neuronal subpopulations in the dorsal root ganglion of the mouse as characterized by combination of ultrastructural and cytochemical features. Brain Res 346(2):310–326.

31. Lawson SN, Harper AA, Harper EI, Garson JA, Anderton BH (1984) A monoclonal antibody against neurofilament protein specifically labels a subpopulation of rat sensory neurones. J Comp Neurol 228(2):263–272.

32. Parysek LM, Goldman RD (1988) Distribution of a novel 57 kDa intermediate filament (IF) protein in the nervous system. J Neurosci 8(2):555–563.

33. Ferri GL, et al. (1990) Neuronal intermediate filaments in rat dorsal root ganglia: differential distribution of peripherin and neurofilament protein immunoreactivity and effect of capsaicin. Brain Res 515(1-2):331–335.34.

34. Bae JY, Kim JH, Cho YS, Mah W, Bae YC (2015) Quantitative analysis of afferents expressing substance P, calcitonin gene-related peptide, isolectin B4, neurofilament 200, and Peripherin in the sensory root of the rat trigeminal ganglion. J Comp Neurol 523(1):126–138.

35. Zhao J, et al. (2010) Small RNAs control sodium channel expression, nociceptor excitability, and pain thresholds. J Neurosci 30(32):10860–10871.

36. Lekan HA, Chung K, Yoon YW, Chung JM, Coggeshall RE (1997) Loss of dorsal root ganglion cells concomitant with dorsal root axon sprouting following segmental nerve lesions. Neuroscience 81(2):527–534.

37. Tandrup T, Woolf CJ, Coggeshall RE (2000) Delayed loss of small dorsal root ganglion cells after transection of the rat sciatic nerve. J Comp Neurol 422(2): 172–180.

38. Spaulding EL, et al. (2016) Synaptic deficits at neuromuscular junctions in two mouse models of Charcot-Marie-Tooth type 2d. J Neurosci 36(11):3254–3267.

39. de Nooij JC, Doobar S, Jessell TM (2013) Etv1 inactivation reveals proprioceptor subclasses that reflect the level of NT3 expression in muscle targets. Neuron 77(6):1055–1068.

40. Alvarez FJ, Morris HR, Priestley JV (1991) Sub-populations of smaller diameter trigeminal primary afferent neurons defined by expression of calcitonin gene-related peptide and the cell surface oligosaccharide recognized by monoclonal antibody LA4. JNeurocytol 20(9):716–731.

41. Scherrer G, et al. (2009) Dissociation of the opioid receptor mechanisms that control mechanical and heat pain. Cell 137(6): 1148–1159.

42. Cavanaugh DJ, et al. (2009) Distinct subsets of unmyelinated primary sensory fibers mediate behavioral responses to noxious thermal and mechanical stimuli. Proc Natl Acad Sci U S A 106(22):9075–9080.

43. McCoy ES, et al. (2013) Peptidergic CGRPα primary sensory neurons encode heat and itch and tonically suppress sensitivity to cold. Neuron 78(1): 138–151.

44. Oliveira Fernandes M, Tourtellotte WG (2015) Egr3-dependent muscle spindle stretch receptor intrafusal muscle fiber differentiation and fusimotor innervation homeostasis. J Neurosci 35(14):5566–5578.

45. Wickramasinghe SR, et al. (2008) Serum response factor mediates NGF-dependent target innervation by embryonic DRG sensory neurons. Neuron 58(4):532–545.

46. Patel TD, Jackman A, Rice FL, Kucera J, Snider WD (2000) Development of sensory neurons in the absence of NGF/TrkA signaling *in vivo*. Neuron 25(2):345–357.

47. Grynkiewicz G, Poenie M, Tsien RY (1985) A new generation of Ca^2+^ indicators with greatly improved fluorescence properties. J Biol Chem 260(6):3440–3450.

48. Caterina MJ, et al. (1997) The capsaicin receptor: a heat-activated ion channel in the pain pathway. Nature 389(6653):816–824.

49. Patapoutian A, Reichardt LF (2001) Trk receptors: mediators of neurotrophin action. Curr Opin Neurobiol 11(3):272–280.

50. Kitao Y, Robertson B, Kudo M, Grant G (1996) Neurogenesis of subpopulations of rat lumbar dorsal root ganglion neurons including neurons projecting to the dorsal column nuclei. J Comp Neurol 371(2):249–257.

51. Maro GS, et al. (2004) Neural crest boundary cap cells constitute a source of neuronal and glial cells of the PNS. Nat Neurosci 7(9):930–938.

52. Stucky CL, DeChiara T, Lindsay RM, Yancopoulos GD, Koltzenburg M (1998) Neurotrophin 4 is required for the survival of a subclass of hair follicle receptors. J Neurosci 18(17):7040–7046.

53. Smeyne RJ, et al. (1994) Severe sensory and sympathetic neuropathies in mice carrying a disrupted Trk/NGF receptor gene. Nature 368(6468):246–249.

54. Ernfors P, Lee KF, Kucera J, Jaenisch R (1994) Lack of neurotrophin-3 leads to deficiencies in the peripheral nervous system and loss of limb proprioceptive afferents. Cell 77(4):503–512.

55. Moqrich A, et al. (2004) Expressing TrkC from the TrkA locus causes a subset of dorsal root ganglia neurons to switch fate. Nat Neurosci 7(8):812–818.

56. Fitzgerald M (2005) The development of nociceptive circuits. Nat Rev Neurosci 6(7):507–520.

57. von Hehn CA, Baron R, Woolf CJ (2012) Deconstructing the neuropathic pain phenotype to reveal neural mechanisms. Neuron 73(4):638–652.

58. Stum M, et al. (2010) An assessment of mechanisms underlying peripheral axonal degeneration caused by aminoacyl-tRNA synthetase mutations. Mol Cell Neurosci 46(2):432–443.

59. Sleigh JN, Schiavo G (2016) Older but not slower: aging does not alter axonal transport dynamics of signalling endosomes *in vivo*. Matters 10.19185/matters.201605000018.

60. Sleigh JN, et al. (2014) Chondrolectin affects cell survival and neuronal outgrowth in *in vitro* and *in vivo* models of spinal muscular atrophy. Hum Mol Genet 23(4):855–869.

61. Sleigh JN, Weir GA, Schiavo G (2016) A simple, step-by-step dissection protocol for the rapid isolation of mouse dorsal root ganglia. BMC Res Notes 9:82.

62. Shepherd AJ, Mohapatra DP (2012) Tissue preparation and immunostaining of mouse sensory nerve fibers innervating skin and limb bones. J Vis Exp (59):e3485.

63. Cesca F, et al. (2012) Kidins220/ARMS mediates the integration of the neurotrophin and VEGF pathways in the vascular and nervous systems. Cell Death Differ 19(2): 194–208.

64. Sleigh JN, Burgess RW, Gillingwater TH, Cader MZ (2014) Morphological analysis of neuromuscular junction development and degeneration in rodent lumbrical muscles. J Neurosci Methods 227:159–165.

65. Vieira JM, Schwarz Q, Ruhrberg C (2007) Selective requirements for NRP1 ligands during neurovascular patterning. Development 134(10): 1833–1843.

65. Wendykier P, Nagy JG (2010) Parallel Colt: A high-performance Java library for scientific computing and image processing. Acm TMath Software 37(3).

66. Dougherty R (2005) Extensions of DAMAS and benefits and limitations of deconvolution in beamforming. 11th AIAA/CEAS Aeroacoustics Conferences, Monterey, California.

67. Nasse MJ, Woehl JC (2010) Realistic modeling of the illumination point spread function in confocal scanning optical microscopy. J Opt Soc Am A 27(2):295–302.

68. Bogdanik LP, et al. (2013) Loss of the E3 ubiquitin ligase LRSAM1 sensitizes peripheral axons to degeneration in a mouse model of Charcot-Marie-Tooth disease. Dis Model Mech 6(3):780–792.

69. Chaplan SR, Bach FW, Pogrel JW, Chung JM, Yaksh TL (1994) Quantitative assessment of tactile allodynia in the rat paw. J Neurosci Methods 53(1):55–63.

70. Carter RJ, Morton J, Dunnett SB (2001) Motor coordination and balance in rodents. Curr Protoc Neurosci / editorial board, Jacqueline N Crawley [et al] Chapter 8, Unit 8 12.

71. Randall LO, Selitto JJ (1957) A method for measurement of analgesic activity on inflamed tissue. Arch Int Pharmacodyn Ther 111(4):409–419.

72. Hargreaves K, Dubner R, Brown F, Flores C, Joris J (1988) A new and sensitive method for measuring thermal nociception in cutaneous hyperalgesia. Pain 32(1):77–88.

73. Domijan AM, Kovac S, Abramov AY (2014) Lipid peroxidation is essential for phospholipase C activity and the inositol-trisphosphate-related Ca^2+^ signal. J Cell Sci 127(Pt 1):21–26.

